# Hippo/YAP1 Signaling Regulates the Oligodendrocyte–Astrocyte Fate Switch and Ependymal Gene Expression in Adult Spinal Cord Stem Cells

**DOI:** 10.1101/2025.10.01.677488

**Authors:** SE Joppe, R Chevreau, J Abou-Chaaya, L Garcia, V Hansberg Pastor, S Hideg, C Ripoll, H Ghazale, A Nouhaud, E Lalli, C Ruggiero, L Zhang, M Chen, S Lugand, S Urbach, K El Koulali, M Seveno, G Poulen, F Vachiery-Lahaye, L Bauchet, JP Hugnot

## Abstract

The adult mammalian spinal cord harbors ependymal cells that retain neural stem-cell properties. Although they possess a latent capacity to generate oligodendrocytes, these cells predominantly differentiate into astrocytes after injury. The molecular cues that govern their lineage commitment toward astrocytic versus oligodendroglial fates remain poorly defined.

In this study, we addressed this gap in vitro by investigating the emergence of PDGFRA⁺ oligodendrocyte precursor cells (OPCs) in neurosphere cultures derived from adult spinal cord stem cells. We first observed that neurosphere cells exhibited a hybrid identity, co-expressing transcription factors of both astrocytic (NFIA, SOX9) and oligodendrocytic (OLIG1/2, SOX4, NKX2.2, TCF4) lineages. Upon differentiation, oligodendrocytic transcription factors were selectively maintained in OPCs but reduced in other cells. Using *Pdgfra*^H2B-GFP^ mice, we then isolated newly formed PDGFRA⁺ OPCs from neurospheres and performed multi-omic profiling. OPC formation was associated with the upregulation of chromatin remodelers and the downregulation of stem-cell markers such as EGFR, HES1, and TNC. Strikingly, OPC specification coincided with reduced expression of YAP1 and its partner TEAD1, key effectors of the Hippo pathway. Functional analyses revealed that YAP1 loss enhanced oligodendrocytic differentiation while reducing astrocytic and ependymal/cilia-associated gene expression. Conversely, constitutive YAP1 activation blocked differentiation into both lineages and promoted an ependymal-like transcriptional program, including upregulation of the ependymal marker CD24a and cilia-related proteins such as CROCC (Rootletin).

Collectively, these findings uncover previously unrecognized roles for YAP1 in adult spinal cord stem-cell fate decisions and provide a molecular framework for leveraging these cells in regenerative strategies targeting spinal cord repair.

## Introduction

In lower vertebrates, the spinal cord retains the capacity to regenerate after injury, owing to a reservoir of ependymal cells that persist around the central canal and resemble fetal radial glia with stem cell properties^1^. Upon spinal cord injury (SCI), these ependymal cells re-enter the cell cycle and give rise to both neurons and glia. Similar ependymal cells are also present in adult mice and humans^1–13^. Organized as an epithelium, they maintain a developmental transcription factor profile—including *Arx*, *FoxA1/2*, *Msx1*, *Meis2*, *Pax6*, *Sox2/4/6/11*—and display embryonic-like dorsoventral patterning, with persistent roof- and floor plate–derived radial cells^3,14,15^. A subset of these ependymal cells can form passageable neurospheres in vitro, a hallmark of neural stem cells^7–10,16–18^. Upon differentiation, these neurosphere cells give rise predominantly to astrocytes, along with a substantial fraction of oligodendrocytes and only a few neurons, confirming their multipotency^9^.

In vivo, murine ependymal cells remain largely quiescent. However, following SCI, they proliferate^19^ and migrate toward the lesion site, where they primarily generate astrocytes and only a few oligodendrocytes^5,13,20–24^, despite exhibiting a chromatin landscape permissive for oligodendrogenesis^25^. Forced expression of the oligodendrocyte lineage-specifying factor OLIG2 is sufficient to redirect these cells toward myelinating oligodendrocytes and promote axonal remyelination after SCI, highlighting their therapeutic potential^25^.

In contrast to rodents, the ependymal region in humans appears to regress after the second decade of life in most individuals^26^. Nevertheless, recent studies have demonstrated that a heterogeneous population of ependymal cells with embryonic features persists throughout adulthood, up to at least 83 years of age^3,11,13,27^. These findings underscore the latent potential of adult human ependymal cells and support efforts to harness them for regenerative therapies.

Although adult spinal cord ependymal cells can give rise to both astrocytic and oligodendrocytic lineages, the molecular cues that govern this fate decision remain poorly understood. In vitro, inflammatory cytokines such as oncostatin M, CNTF, IFN-γ, and TNF-α, along with BMP signaling, promote astrocytic differentiation while repressing oligodendrocyte-associated markers^9,21^. However, additional signaling pathways likely contribute to this lineage specification. Furthermore, oligodendrocyte formation typically occurs via intermediate oligodendrocyte progenitor cells (OPCs), which are characterized by specific expression of the PDGFRA receptor^28^. It remains unclear whether adult spinal cord ependymal cells can produce PDGFRA⁺ OPCs and, if so, through which mechanisms.

In this study, we sought to address these gaps by investigating whether adult spinal cord stem cells can generate OPCs and elucidating the mechanisms underlying this process. To this end, we used neurosphere cultures derived from the adult mouse spinal cord, which we and others have shown originate primarily from ependymal cells^10,22^. By leveraging neurospheres from adult *Pdgfra*^H2B-GFP^ knock-in mice^29^, we were able to isolate newly formed PDGFRA⁺ cells and profile them using RNA sequencing and quantitative proteomics. Combined with gain- and loss-of-function approaches, these analyses identified the Hippo/YAP1 signaling pathway as a regulator of spinal cord stem cell proliferation, commitment into astrocytic vs oligodendrocytic lineages and the expression of ependymal and cilia-associated genes.

## Results

### Adult spinal cord neurospheres express transcription factors associated with both oligodendrocyte and astrocyte lineages

To investigate the emergence of oligodendrocyte lineage cells from mouse spinal cord ependymal cells, we generated neurosphere cultures from adult mice and examined by immunofluorescence (IF) six key oligodendrocyte transcription factors (NKX2.2, OLIG1/2, SOX4, SOX10, TCF4)^30^. Under proliferative conditions (with EGF/FGF2), most neurosphere cells expressed these factors at variable levels, except SOX10, which was undetectable by IF or WB despite a validated antibody (Fig. 1A; Fig. S1A,B). The two astrocytic transcription factors^31^ NFIA and SOX9 were also broadly expressed in neurosphere cells (Fig. 1B). These findings indicate that proliferating spinal cord stem cell cultures constitute a hybrid population co-expressing both astrocytic and oligodendrocytic transcription factors with marked cell-to-cell variability.

**Figure 1.**
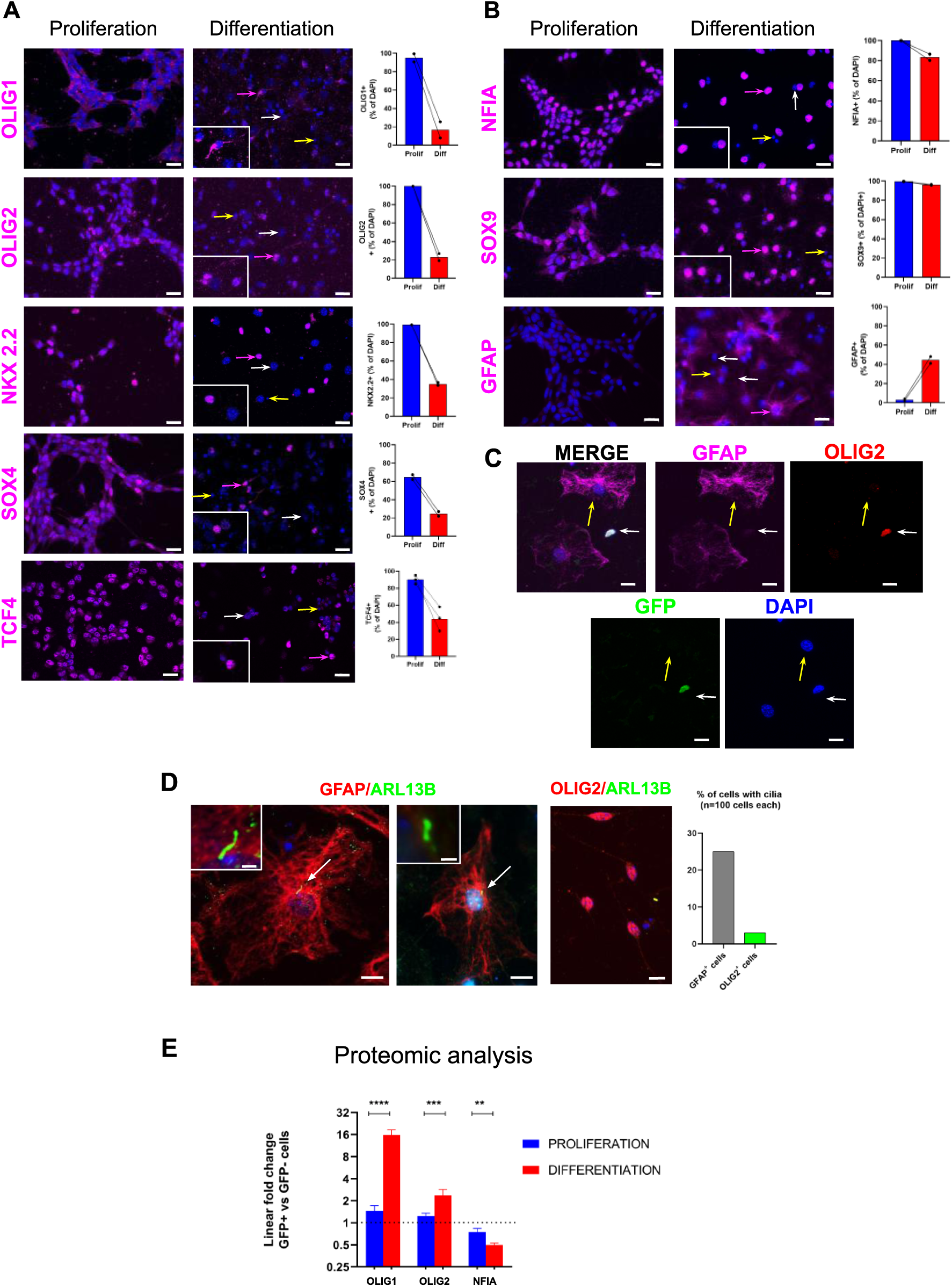
**Regulation of oligodendrogenic and astroglial transcription factors in adult spinal cord neurospheres.** A. Immunofluorescence for oligodendrocyte transcription factors (TF) (OLIG1, OLIG2, NKX2-2, SOX4, TCF4) in neurospheres cultured under proliferative conditions (EGF + FGF2) or after 5 days without growth factors (differentiation). White arrows indicate TF-negative nuclei, pink arrows TF-positive nuclei, and yellow arrows dead cells. Graphs show the mean ± SEM percentage of TF⁺ cells from two independent cultures (three for TCF4). These TFs are broadly expressed during proliferation but markedly reduced upon differentiation. Scale: 20 µm. B. Equivalent analysis for astroglial TFs (NFIA, SOX9) and GFAP. NFIA and SOX9 are maintained, while GFAP increases upon differentiation. Scale: 20 µm. C. Differentiated PDGFRA-GFP⁺ neurospheres stained for GFAP (magenta), OLIG2 (red), GFP (green). Distinct astrocytic (GFAP⁺, yellow arrows, round nucleus) and oligodendroglial (OLIG2⁺/GFP⁺, white arrows, small, oval nucleus) populations are evident. Scale: 10 µm D. Primary cilia analysis using ARL13B (green) in differentiated neurospheres. GFAP⁺ cells show ARL13B⁺ cilia (arrows, insets), while OLIG2⁺ cells largely lack them. Quantification (right) shows the proportion of ciliated cells (n = 100 cells per group). Scale: 10 µm; insets: 1 µm. E. Proteomics of OLIG1, OLIG2, NFIA in PDGFRA-GFP⁺ vs GFP⁻ cells under proliferation (blue) and differentiation (red). Values are linear fold-change (GFP⁺/GFP⁻); paired t-test (**p<0.01, ***p<0.001, ****p<0.0001). Nuclei, DAPI (blue), all photographs.

Following a five-day growth factor withdrawal to promote differentiation, SOX10 remained absent (Fig. S1A,B), whereas OLIG1/2, NKX2.2, SOX4, and TCF4 expression declined sharply and became restricted to a subset of bipolar cells (e.g., inset OLIG1 staining, Fig. 1A) with small, elongated nuclei (Fig. 1A,C,D). In contrast, a second major population— larger polygonal cells with pale, round nuclei—displayed little or no expression of these factors but expressed GFAP at variable levels, consistent with astrocytic identity (Fig. 1B–D). Approximately 25% of these astrocytic cells exhibited a primary cilium (detected by Arl13b⁺), suggestive of quiescence, whereas cilia were rarely observed in OLIG2⁺ cells (Fig. 1D). NFIA and SOX9 showed only a moderate reduction and remained variably expressed in both populations (Fig. 1B).

We next assessed the timing of OPC formation in this spinal cord neurosphere model by monitoring PDGFRA, a specific OPC marker^28^. WB revealed low PDGFRA expression under proliferative conditions, but a marked increase upon differentiation. Consistently, IF showed that PDGFRA⁺ cells were rare in proliferating cultures but accounted for ∼5–10% of cells after differentiation (Fig. 2B). These cells were characterized by smaller nuclei than neighboring cells and a bipolar or multibranched morphology (Fig. 2B, D), consistent with oligodendrocyte lineage identity.

**Figure 2.**
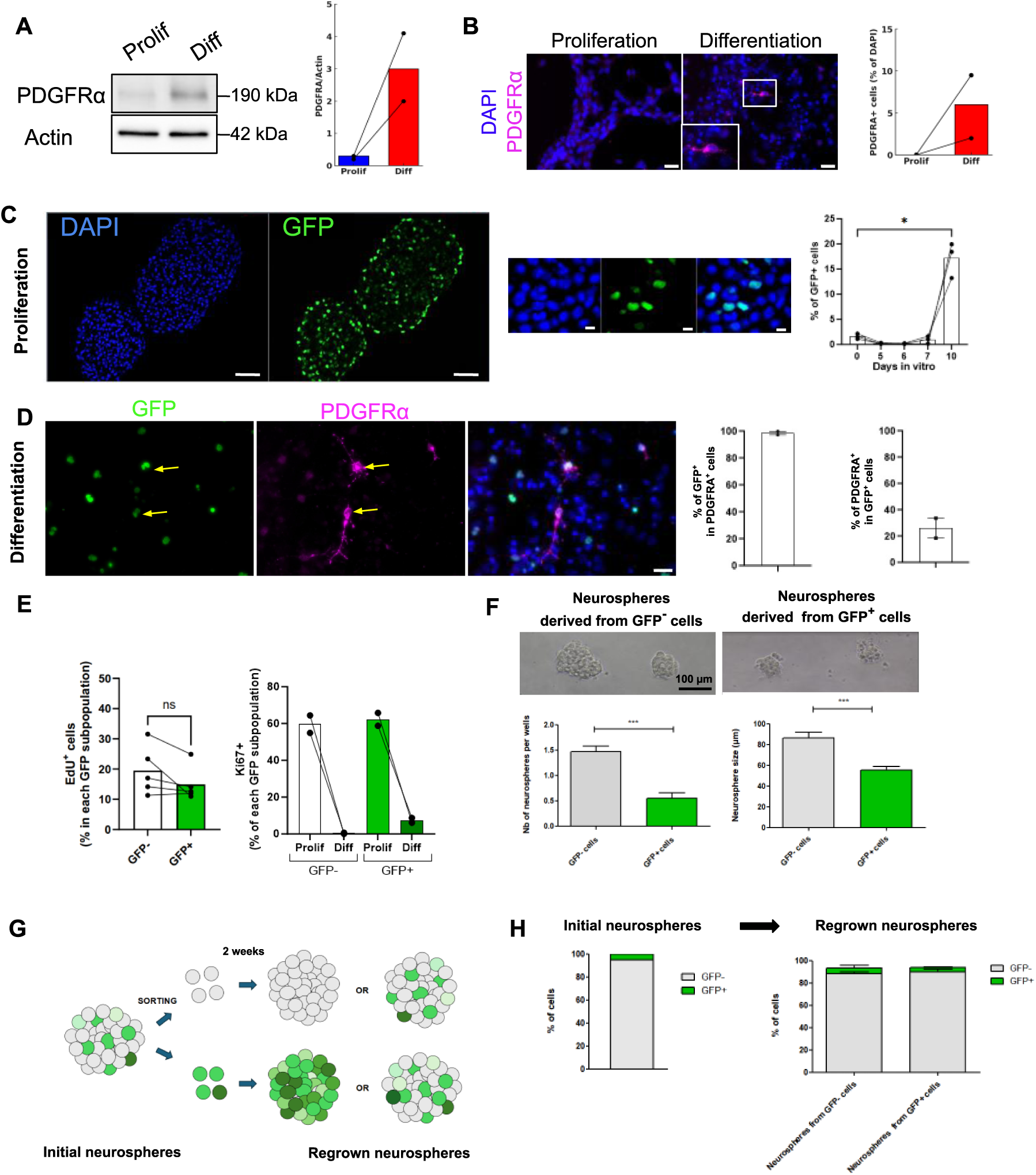
**PDGFRA expression, proliferation, and plasticity of PDGFRA-GFP⁺ neurosphere cells.** A. Western blot of PDGFRA in proliferating (Prolif) vs. differentiating (Diff, 5 days) cultures, with Actin as control. Quantification (right, n = 2) shows increased PDGFRα after differentiation. B. Immunofluorescence for PDGFRA (magenta) under Prolif vs. Diff conditions. Inset: typical bipolar morphology of a PDGFRA-GFP⁺ OPC cell. Quantification confirms increased PDGFRα⁺ cells (n = 2). Scale: 20 µm. C. Time-course of GFP expression in neurospheres derived from PDGFRA^-H2B-GFP^ mice. Left: neurosphere section (10 days in culture) showing GFP⁺ cells. scale: 40 µm. Middle: magnified view (scale: 10 µm). Right: FACS quantification of GFP⁺ cells over time (n = 3 independent cultures). GFP⁺ cells appear abruptly after ∼10 days in culture. test: two-tailed t-test; p < 0.05 ().* D. Co-staining for PDGFRA and GFP after neurosphere differentiation. Yellow arrows: double-positive cells. Right: quantification of double-positive cells among the total PDGFRα⁺ or GFP⁺ populations (mean ± s.e.m., n=2 experiments). E. Proliferation of GFP⁺ and GFP⁻ cells. EdU incorporation (left, n = 4; ns: not significant) and Ki67 staining (right, n = 2) show similar proliferation rates in both populations under proliferative conditions. F. Neurosphere formation from GFP⁺ and GFP⁻ cells. FACS-sorted cells (10 per well, 96-well plate) were cultured for 2 weeks. GFP⁺ cells generated fewer and smaller neurospheres than GFP⁻ cells (top: representative images, scale bar: 100 µm; bottom: quantification from three independent experiments; ***p < 0.001, two-tailed t-test). G. Experimental design to assess GFP⁺ cell plasticity. Sorted GFP⁺ and GFP⁻ cells were cultured separately to form neurospheres. After 2 weeks, neurospheres were dissociated and re-analyzed by FACS to determine whether each population remained stable or could regenerate the other. H. Flow cytometry of GFP⁺ and GFP⁻ proportions in initial and regrown neurospheres. After 2 weeks, neurospheres derived from either GFP⁺ or GFP⁻ cells were composed predominantly of GFP⁻ cells with a minority of GFP⁺ cells (n = 3 experiments). G. Experimental design: sorted GFP⁺ and GFP⁻ cells cultured separately, then re-analyzed after 2 weeks to assess plasticity. H. Flow cytometry of regrown neurospheres shows both sorted populations regenerate mixed neurospheres, predominantly GFP⁻ with a minority of GFP⁺ cells (*n* = 3).

To further track PDGFRA⁺ cell emergence, we generated neurosphere cultures from heterozygote *Pdgfra*^H2B-GFP^ knock-in mice. Appearance of PDGFRA-GFP⁺ cells, hereafter referred to as GFP⁺ cells, were monitored over time by cytometry. As shown in Figure 2C, GFP⁺ cells appeared abruptly around day 10 of culture, coinciding with the formation of large neurospheres (>200 µm). During proliferation, PDGFRA protein in GFP⁺ cells was very low or undetectable by IF (data not shown), suggesting they represent an early stage of commitment toward the OPC state. In contrast, growth factor withdrawal led to robust PDGFRA expression in ∼30% of GFP⁺ cells (Fig. 2D), the majority of which (>90%) co-expressed OLIG2 (Fig. S2B). The remaining GFP⁺ cells were PDGFRA⁻, consistent with an immature state lacking sufficient receptor expression. Importantly, all PDGFRA⁺ cells were GFP⁺, confirming the specificity of the reporter (Fig. 2D). These findings highlight both the ability of spinal cord neurospheres to generate PDGFRA⁺ subpopulations and the dynamic regulation of PDGFRA during differentiation, underscoring the value of *Pdgfra*^H2B-GFP^ mice for identifying and isolating these cells. We further characterized PDGFRA-GFP⁺ and GFP⁻ cells by IF. Under proliferative conditions, both populations expressed comparable levels of oligodendrocyte regulators (NKX2.2, OLIG1/2, SOX4) and astrocyte regulators (NFIA, SOX9) (Fig. S2A). Upon differentiation, however, GFP⁻ cells lost oligodendroglial determinants, whereas GFP⁺ cells retained them, downregulated NFIA, and maintained SOX9 (Fig. S2B). Proteomic analyses of both populations (see below) mirrored these changes: OLIG1, OLIG2, and NFIA were similar in both populations under proliferative conditions but diverged after differentiation, with oligodendroglial regulators enriched in PDGFRA-GFP⁺ cells (Fig. 1E).

Together, these findings support a model in which adult spinal-cord neurospheres contain cells that co-express astrocytic and oligodendrocytic transcription factors, and that oligodendrocyte lineage commitment is driven by the selective retention—or loss—of these factors during differentiation.

### Transcriptomic and proteomic analyses confirm that PDGFRA-GFP⁺ cells belong to the oligodendrocyte lineage

To characterize PDGFRA-GFP⁺ cells in an unbiased manner, we FACS-sorted them from neurospheres and compared their RNA profiles with GFP⁻ cells (Table S1). Under proliferative conditions, 582 genes were enriched in PDGFRA-GFP⁺ cells and 1379 in GFP⁻ cells (|log₂ fold change| >1, q<0.05) (Fig. 3A). Gene Set Enrichment Analysis (GSEA)^32^ confirmed that PDGFRA-GFP⁺ cells are enriched in OPC and oligodendrocyte gene signatures, including *Ascl1, Cspg4, Myt1, Olig1/2, Pdgfra*, and *Sox8* (Fig. 3B,C). In contrast, GFP⁻ cells were enriched for ependymal, astrocytic (*Cryab, Id3, Nfia, Pax6, Sparc*), and radial glial markers (*Fabp7, Hopx, Slc1a3, Vim*), reflecting their broader developmental potential (Fig. 3B,C).

**Figure 3.**
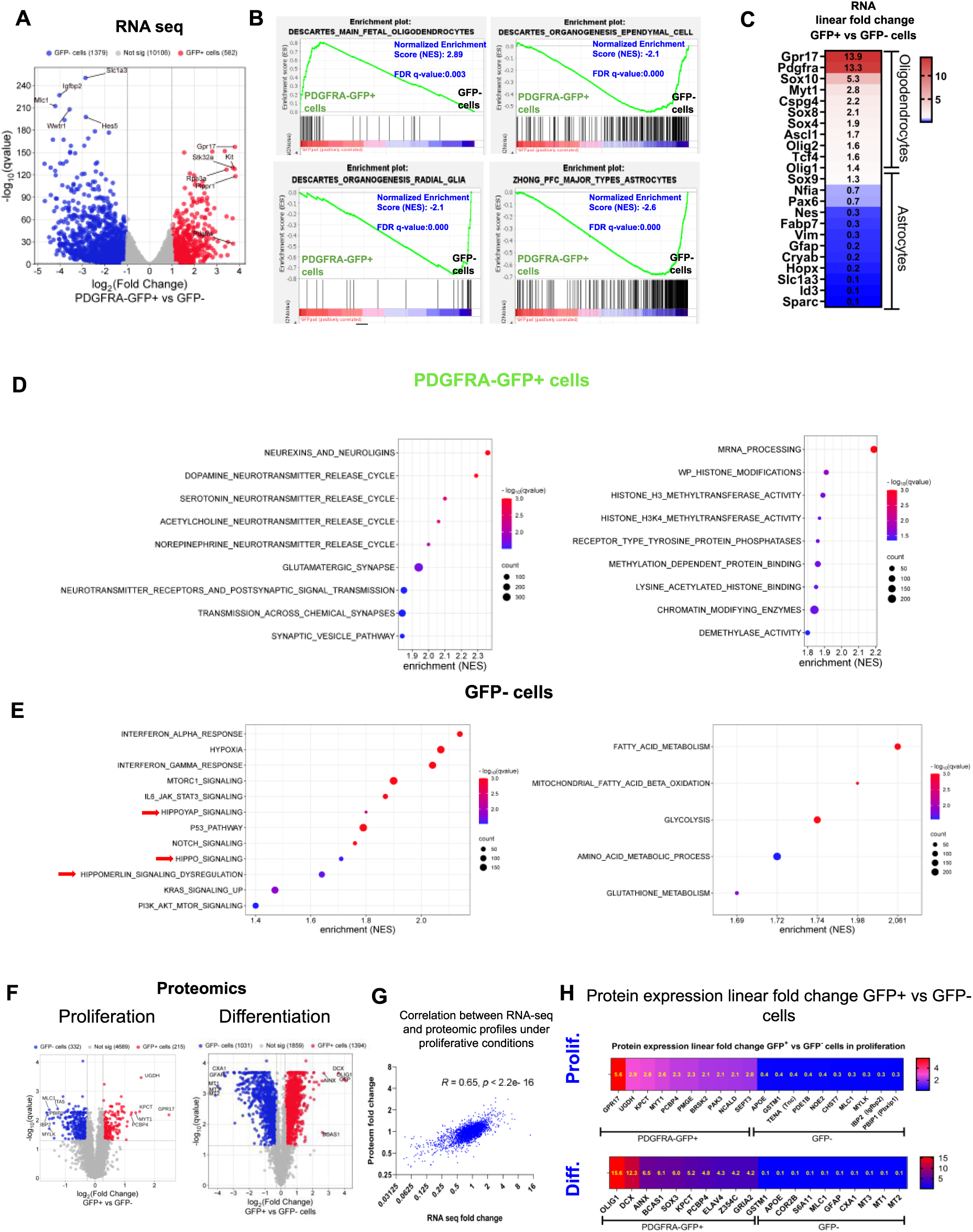
**Transcriptomic and proteomic profiles of PDGFRA-GFP⁺ and GFP⁻ spinal cord neurosphere cells** A. Volcano plot of bulk RNA-seq comparing GFP⁺ and GFP⁻ cells under proliferative conditions (n = 3). Differentially expressed genes were defined as those with log_2_(|fold change| > 1 and q-value < 0.05. B. GSEA (Gene Set Enrichment Analysis) of bulk RNA-seq. Using an established OPC gene list, GFP⁺ cells show strong enrichment with reciprocal depletion in GFP⁻ cells, whereas GFP⁻ cells are enriched for radial glia, astrocyte, and ependymal markers. C. Heatmap (linear fold change) of selected differentially expressed genes between GFP⁺ vs GFP⁻ cells (p < 0.05). GFP⁺ cells show enriched expression of OPC markers (e.g., *Gpr17, Pdgfra, Cspg4, Ascl1, Olig2)*, while GFP⁻ cells are enriched for astrocytic markers (e.g., *Gfap, Hopx, Sparc, Fabp7*) D–E. Bubble charts of selected pathways from Table S1 significantly enriched in GFP⁺ and GFP⁻ cells. Red arrows in panel E highlight enrichment of Hippo/YAP1 signaling components in GFP⁻ cells. F. Volcano plots of proteomic profiling of GFP⁺ versus GFP⁻ cells under proliferative and differentiative conditions. Differentially expressed proteins were defined using thresholds of |linear fold change| > 1.2 (i.e., |log₂ fold change| > 0.26) and q-value < 0.05 (n = 5). G. Scatter plot showing correlation between RNA-seq and proteomic fold changes, demonstrating positive concordance between datasets (Pearson r = 0.65, p < 0.001). H. Heatmap (linear fold change) of selected differentially expressed proteins (p < 0.05) between GFP⁺ and GFP⁻ cells. Proteins representative of the oligodendrocyte lineage are shown in red/purple and those of the astrocytic lineage in blue. Linear fold changes (GFP⁺ vs GFP⁻) are indicated in yellow for each protein.

Functionally, PDGFRA-GFP⁺ cells showed pronounced enrichment of synaptic signaling and neurotransmission pathways (Fig. 3D); glutamate-receptor subunits Grin1, *Grin3*, and *Grik3* increased ∼4.4-, 5.3-, and 7-fold, and the scaffold *Shank1* ∼4.7-fold (Table S1), aligning with reports that OPCs engage in neuron–glia synapses^33^. They also displayed elevated RNA-processing and chromatin-modification programs (Fig. 3D); for example, *Chd3* and *Prmt8* rose ∼4.8- and 6.9-fold, suggesting active chromatin reorganization during OPC specification. Conversely, GFP⁻ cells were enriched for metabolic and stem-cell–associated signaling (PI3K–AKT–mTOR, Notch) and showed higher Hippo/YAP1 activity (Fig. 3E), consistent with down-modulation of this pathway in emerging PDGFRA-GFP⁺ cells. GFP⁻ cells also displayed interferon-response gene enrichment. A Cytoscape^34^ summary of GSEA results (Fig. S2C) recapitulated these contrasts: synaptic/RNA–DNA processing in PDGFRA-GFP⁺ cells versus ECM/collagen, metabolism, energy production, and stem-cell programs in GFP⁻ cells.

Proteomic profiling further confirmed these results. Among the 5,237 proteins detected, 215 were enriched in PDGFRA-GFP⁺ cells and 332 in GFP⁻ cells (|linear fold change| >1.2, q<0.05; Fig. 3F, Table S1). Transcriptomic and proteomic datasets showed strong concordance (Pearson r=0.65, p<0.001; Fig. 3G). The top PDGFRA-GFP⁺-specific proteins (Fig. 3H, upper panel) were predominantly associated with the oligodendrocyte lineage, as revealed by over-representation analysis^35^ (ORA), GSEA^32^ (Table S1), and adult spinal cord single-cell atlases^36^. These included MYT1 and GPR17, well-known regulators of oligodendrogenesis, as well as novel candidates such as BRSK2, NCALD, SEPT3, and UGDH. PDGFRA-GFP⁺ cells also displayed elevated expression of proteins involved in chromatin remodeling (KDM4A/B, KDM5B, JMJD1C) and neuron–OPC interactions (GRIA2) (Table S1). By contrast, GFP⁻ cells were enriched for astrocytic proteins (APOE, SLC1A3, SPARC, TNC) and markers of ependymal/astrocytic identity (IBP2, PBXIP1, GSTM1, MLC1) as identified in an adult spinal cord single-cell atlas (Fig. 3H, Fig. S4)^36^.

Finally, under differentiation conditions (i.e. w/o growth factors), PDGFRA-GFP⁺ cells remained enriched in oligodendrocyte-associated proteins (1394 upregulated), while GFP⁻ cells preferentially expressed astrocytic proteins (1031 upregulated) (|linear fold change| >1.2, q-value<0,05, Fig. 3F,H lower panel). Growth factor withdrawal induced strong astrocytic signatures in GFP⁻ cells, including AQP4, CXA1 (*Gja1*), GFAP, and SPRL1 (*Sparcl1*), which were absent under proliferation (Table S1).

Together, these transcriptomic and proteomic data consistently shows that PDGFRA-GFP⁺ cells formed in neurosphere are committed to the oligodendrocyte lineage, whereas GFP⁻ cells display astrocytic, radial glial, and ependymal features.

### PDGFRA-GFP⁺ cells exhibit reduced neurosphere-forming ability and can revert to a GFP⁻ state

Because PDGFRA-GFP⁺ cells display distinct molecular profiles, we next examined their functional properties. Proliferation analysis using EdU incorporation and KI67 staining revealed no significant difference between PDGFRA-GFP⁺ and GFP⁻ cells, with ∼60% of each population actively cycling under proliferative conditions (Fig. 2E). Growth factor withdrawal for five days reduced KI67 expression similarly in both populations (Fig. 2E).

In contrast, low-density neurosphere assays (0.1 cells/µl, a common method to assess stem cell properties^37^) revealed that PDGFRA-GFP⁺ cells exhibited a reduced ability to form neurospheres, which were also smaller on average than those derived from GFP⁻ cells (Fig. 2F). To test cell plasticity, neurospheres derived from sorted PDGFRA-GFP⁺ or GFP⁻ cells were expanded for two weeks and analyzed by cytometry. Strikingly, most cells from PDGFRA-GFP⁺-derived neurospheres lost GFP expression, resembling the phenotype of GFP⁻-derived neurospheres (Fig. 2G,H).

These findings indicate that while PDGFRA-GFP⁺ cells form fewer and smaller neurospheres, at least a fraction of them are not irreversibly committed to this state and can revert to a GFP⁻ phenotype, underscoring their intrinsic plasticity.

### The formation of PDGFRA-GFP+ cells is associated with the downregulation of proteins involved in Ras, Notch1 and Hippo/YAP1 pathways

Pathway analysis showed that, compared to GFP⁻ cells, PDGFRA-GFP⁺ cells exhibited reduced mTOR/RAS and Notch signaling (Fig. 3E). Consistently, IF, RNA-seq, and proteomics revealed strong downregulation of EGFR, TNC, and HES1—key regulators of these pathways and of neural stem cell maintenance (Fig. 4A,B). RNA-seq further showed a marked decrease in Notch transcriptional targets (*Hey1, Hey2, Hes1, Hes5*) (Fig. 4B), which were undetectable by proteomics, likely due to low abundance.

**Figure 4.**
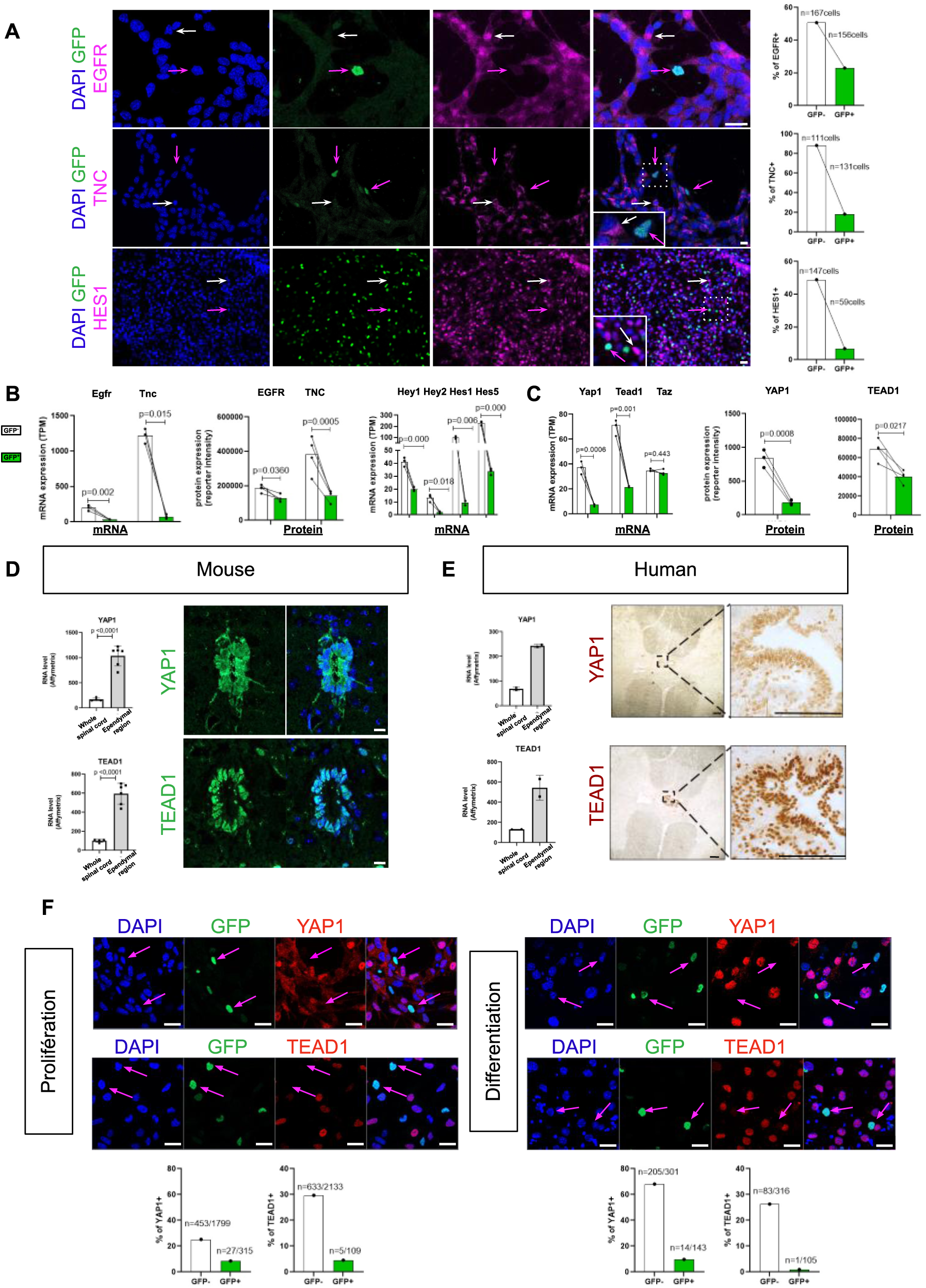
**Formation of PDGFR-GFP⁺ cells is accompanied by reduced expression of EGFR, TNC, Notch-regulated transcription factors, YAP1, and TEAD1** A. IF for EGFR, TNC, and HES1 (magenta) in PDGFRA-GFP⁺ spinal cord stem cells under proliferative conditions. Neurospheres were plated in adhesion for 24 h before staining. White arrows indicate GFP⁻ cells, pink arrows GFP⁺ cells. Insets show higher-magnification views of boxed regions. Bar graphs (right) quantify the percentage of marker-positive cells in GFP⁻ and GFP⁺ populations, with cell numbers indicated above each bar. Scale: 10 μm. B. Left: Bar graphs showing mean mRNA (RNA-seq, n = 3) and protein (proteomics, n = 4) expression of EGFR/*Egfr* and TNC/*Tnc* in FACS-sorted PDGFRA-GFP⁺ (green) and GFP⁻ (white) spinal cord cells, confirming the IF in Figure 4A. Right: Mean mRNA expression (RNA-seq, n = 3) of four Notch-regulated transcription factors (*Hey1, Hey2, Hes1, Hes5*) in PDGFRA-GFP⁺ and GFP⁻ cells, consistent with reduced Notch signaling in GFP⁺ cells and supporting the HES1 immunofluorescence in Figure 4A. Tests: two-tailed t-test. C. Bar graphs showing mean mRNA (RNA-seq, n = 3) and protein (n = 3–4) expression of core Hippo pathway components (*Yap1, Tead1, Taz*) in PDGFRA-GFP⁺ (green) and GFP⁻ (white) cells. GFP⁻ cells show significantly higher YAP1/*Yap1* and TEAD1/*Tead1* expression, while *Taz* levels are comparable. Tests: two-tailed t-test. D–E. Expression of YAP1 and TEAD1 in mouse and human spinal cord ependymal cells. Left: Bar graphs showing mean mRNA expression of Yap1/Tead1 in the microdissected ependymal region versus remaining spinal cord tissue, in adult mice (n = 4) and in two human donors (ages 17 and 42), based on Affymetrix microarray data (Ghazale et al., 2019). Tests: two-tailed t-test. Right: Immunofluorescence (mouse) and IHC (human, 53 years) showing YAP1 and TEAD1 expression in ependymal cells. Magnified views highlight the boxed regions. Scale: mouse, 10 µm; human, 200 µm. F. Immunofluorescence for GFP (green), TEAD1 and YAP1 (red) in adult spinal cord neurospheres derived from PdgfraH2B-GFP mice under proliferative and differentiating conditions. For proliferative conditions, neurospheres were plated in adhesion for 24 h before staining. Pink arrows indicate GFP⁺ cells lacking detectable YAP1 or TEAD1. Scale: 10 μm. Quantification of YAP1⁺ and TEAD1⁺ cells within GFP⁺ and GFP⁻ populations is shown below each condition, with marker-positive cell counts indicated above each bar. Nuclei, DAPI (blue), all photographs.

In silico predictions also pointed to reduced Hippo/YAP1 pathway activity in PDGFRA-GFP⁺ cells (Fig. 3E), an intriguing result since this pathway is a central regulator of stem cell maintenance across multiple systems^38^ but remains largely unexplored in OPC formation and adult spinal cord stem cells. Our previous transcriptomic analyses of human and mouse ependymal cells^15^ revealed strong enrichment of Hippo/YAP1 components. Consistently, TEAD1 and YAP1—the pathway’s key transcriptional effectors—were highly expressed in ependymal cells at both RNA and protein levels (Fig. 4D,E). Single-cell RNA-seq atlases of mouse and human spinal cord^36,39,40^ further confirmed elevated YAP1 expression in ependymal cells compared to oligodendrocyte-lineage populations (Fig. S5A,B).

These observations prompted us to assess Hippo/YAP1 expression and function during PDGFRA⁺ cell formation in neurospheres. Comparative RNA-seq and proteomics analyses revealed a significant reduction of YAP1 and TEAD1 in PDGFRA-GFP⁺ cells relative to GFP⁻ cells (Fig. 4C), along with decreased expression of canonical Hippo/YAP1 gene targets *Ccn1/Cyr61* and *Ccn2/Ctgf* (Fig. S5C). In contrast, *Taz* (*Wwtr1*), another Hippo coactivator, showed no change between the two populations (Fig. 4C). IF analysis of neurosphere cultures under both proliferative and differentiated conditions confirmed a marked reduction of YAP1 and TEAD1 protein levels in PDGFRA-GFP⁺ cells (Fig. 4F).

Together, these findings demonstrate that OPC emergence involves coordinated suppression of stem-cell maintenance pathways—RAS/mTOR, Notch1, and Hippo/YAP1— accompanying oligodendrocyte lineage commitment.

### Loss of YAP1 reduces the expression of astrocytic, ependymal-cell, and cilia-related genes, while up-regulating genes associated with the oligodendrocyte lineage

Because YAP1 is enriched in GFP⁻ cells but diminished in PDGFRA-GFP⁺ cells, we hypothesized that it sustains spinal cord stem cells and modulates OPC formation. To test this, we generated neurospheres from *Yap1*^fl/fl^ ^41,42^ and wild-type (wt) mice and ablated *Yap1* through Cre-mediated excision of exon 2 delivered by a viral vector. WB and IF confirmed complete protein loss (Fig. 5A,B). Despite YAP1’s known role in stem-cell proliferation, knockout (KO) neurospheres showed no reduction in growth: EdU incorporation, Ki67 staining, RNA expression of canonical proliferation genes (*Mki67, Pcna, Top2a, Ccnb1*) were unchanged (Fig. 5C, Fig. S6A), and KO cells could be expanded for at least 3 passages without observing any notable disparities in cell expansion.

**Figure 5.**
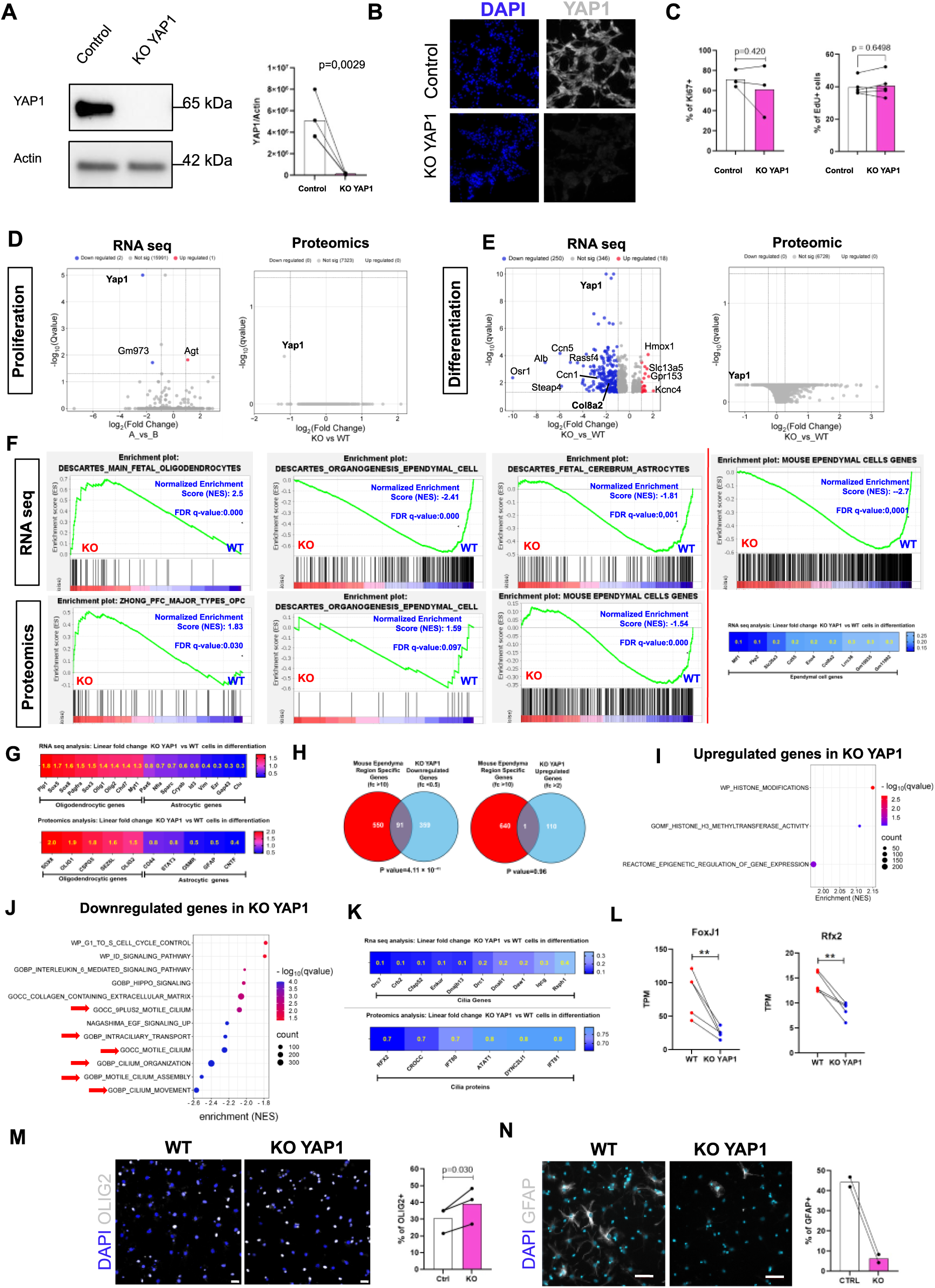
**Analysis of YAP1 KO adult spinal cord neurospheres** A. WB showing complete loss of YAP1 in Yap1^fl/fl^ spinal cord neurospheres treated with Adenovirus-Cre hereafter referred to as YAP1 knockout (KO). Left: Blot for YAP1 and Actin in control and Cre-treated Yap1^fl/fl^-derived spinal cord neurospheres; YAP1 is undetectable in Cre-treated samples using a C-terminal antibody, confirming efficient recombination. Right: Quantification of YAP1/Actin signal intensity (unpaired t-test, n = 3). B. IF for YAP1 (grayscale) in control and YAP1 KO spinal cord neurospheres, confirming loss of YAP1 in KO cells. Scale: 10 µm. C. Quantification of Ki67⁺ (left) and EdU⁺ (right) cells in control and YAP1 KO neurospheres. No significant differences were detected (n = 3 for Ki67, n = 4 for EdU; paired two-tailed t-test; p values shown). D–E. Volcano plots of bulk RNA-seq and proteomics comparing control and YAP1 KO neurospheres under proliferative (D) and differentiating (E) conditions. Differential expression was defined as |log_2_(fold change)| > 1 with q-value < 0.05. In proliferative conditions (D), few genes were significantly up-(red) or downregulated (blue), and no proteins met significance criteria. In differentiating conditions (E), more genes were differentially expressed, indicating a stronger transcriptional impact of YAP1 loss, while proteomic changes remained below significance thresholds. Notably, although YAP1 protein is undetectable by WB and IF (Panels A–B), partial detection by mass spectrometry likely reflects N-terminal fragments upstream of floxed exon 2. F. GSEA of cell type–specific signatures in RNA-seq (top) and proteomics (bottom) from WT and YAP1 KO differentiated neurospheres. KO cells show enrichment of oligodendrocyte lineage signatures and significant depletion of ependymal and astrocyte signatures. NES and FDR q-values are indicated. The bottom-right heatmap shows nine ependymal-enriched genes (see Fig. S8), not directly linked to cilia structure, that are strongly downregulated in KO cells under differentiation. G. Heatmaps of selected differentially expressed genes (top) and proteins (bottom) (p < 0.05) in YAP1 KO versus WT neurospheres under differentiation. Markers of the oligodendrocyte lineage (left, red) and astrocytic lineage (right, blue) are shown, with linear fold changes (KO vs WT) indicated in yellow. H. Venn diagrams showing overlap between mouse ependymal genes (fold change >10; gene list (Table S4) derived from Table S1 in Ghazale et al., *Stem Cell Reports*, 2019) and genes downregulated (fc < 0.5, left) or upregulated (fc > 2, right) in differentiated YAP1 KO neurospheres. Downregulated genes show strong enrichment (hypergeometric test, p = 4.11 × 10⁻⁴¹), whereas no enrichment is observed among upregulated genes (p = 0.96), indicating that YAP1 loss reduces expression of a subset of ependymal genes. I–J. Bubble charts of selected pathways from Table S2 significantly enriched in YAP1 KO and WT cells. K. Heatmaps of 10 cilia-related genes (top) and 6 proteins (bottom) significantly downregulated (p < 0.05) in YAP1 KO neurospheres under differentiation. Linear fold changes (KO vs WT) are indicated in yellow within the boxes.. L. RNA-seq expression of Foxj1 and Rfx2, two key ciliogenesis transcription factors, in WT and YAP1 KO cells (TPM, n = 4; paired t-test, p < 0.01). M–N. Left: Representative IF images of OLIG2 and GFAP (grayscale) in WT and YAP1 KO neurospheres under differentiation. Scale: 10 µm (OLIG2) and 50 µm (GFAP). Right: Quantification of OLIG2⁺ (n = 3, paired t-test) and GFAP⁺ cells (n = 2). YAP1 loss significantly increases OLIG2⁺ cells and reduces GFAP⁺ cells. Nuclei, DAPI (blue), all photographs.

To further assess the effects of YAP1 loss, we profiled KO and wt neurospheres by RNA-seq and proteomics under proliferative and differentiation (Fig. 5D; Table S2, n=5). Under proliferative conditions, changes were minimal, with only three transcripts significantly altered—*Yap1, Gm973*, and the astrocytic marker *Agt* (|log₂ fold change| > 1, q < 0.05; Table S2).

In contrast, YAP1 loss had a pronounced impact during differentiation. RNA-seq revealed 18 upregulated and 250 downregulated genes (|log₂ fold change| > 1, q < 0.05; Fig. 5E), including strong reduction of the Hippo/YAP1 target *Ccn1/Cyr61*. GSEA and ORA showed enrichment of oligodendrocyte lineage genes and regulators of histone modification among upregulated transcripts (Fig. 5F,I), whereas downregulated genes were enriched for astrocytic and ependymal markers (Fig. 5F; Table S2). A heatmap illustrates this reciprocal regulation of oligodendrocytic and astrocytic genes (Fig. 5G, upper panel). To evaluate how YAP1 loss specifically impacts ependymal-cell genes, we used our previously generated human and mouse spinal cord ependymal-cell RNA profiles^15^ to define two gene sets containing transcripts enriched more than tenfold in these cells (Table S4). Applying GSEA to these 2 gene sets revealed a marked depletion of ependymal markers following YAP1 loss (Fig. 5F, far right; Fig. S6C). Venn analyses showed highly significant overlap between KO YAP1 downregulated genes and ependymal-specific transcripts—91 in mice (Fig. 5H; Table S5) and 26 in human datasets (Fig. S6D)—while upregulated genes showed no such enrichment. The heatmap in Fig. 5F (far right) highlights a subset of nine genes showing marked downregulation in KO YAP1 cells (fold change >3), all of which are almost exclusively expressed in ependymal cells according to the single-cell spinal cord atlas (Fig. S7).

Pathway analyses also revealed that YAP1 loss downregulated genes involved in cell cycle regulation and extracellular matrix organization, particularly collagens (Fig. 5J). Collagen type VIII α2 (*Col8a2*), highly specific to ependymal cells according to single-cell atlases (Fig. S7), decreased nearly five-fold in KO neurospheres (Table S2). Unexpectedly, ciliogenesis genes were also strongly reduced (Fig. 5J), including the cilia-transcriptional master regulators *Rfx2*^43^ and *FoxJ1*^44^ (Fig. 5L) and structural components^45^ such as *Bbs1, Cep128, Dnaaf5, Dnal1, Dnali1, Ift122, Ift80, Tctn2*, and *Ttc26* (Table S2). A heatmap highlights the most affected cilia-related genes in KO cells (Fig. 5K). Further evidence supporting YAP1’s role in ciliogenesis comes from the 91 ependymal-specific transcripts that were significantly reduced upon YAP1 deletion, as identified by our Venn analysis (Fig. 5H; Table S5). Over-representation analysis (ORA) of these transcripts revealed strong enrichment for ciliogenesis pathways (Fig. S6E).

Proteomic profiling during differentiation (n=5) was consistent with RNA-seq findings. Although variability between samples prevented significance at q < 0.05, relaxing the threshold (p < 0.05) identified 222 differentially expressed proteins (|fold change| > 1.2) (Table S2; Fig. S6F). Upregulated proteins in KO YAP1 cells included CSPG5, OLIG1, OLIG2, SEZ6L, and SOX8, all linked to the oligodendrocyte lineage (Fig. 5G; Fig. S6F), whereas downregulated proteins included astrocytic markers (GFAP, CD44) and regulators of astrocyte formation (OSMR, CNTF, STAT3) (Fig. 5G, lower panel). Proteomic data also confirmed reduced expression of ciliogenesis proteins, notably the regulator RFX2^44^,as shown in the heatmap in Fig. 5K.

The convergence of our RNA-seq and proteomic datasets supports YAP1 as a key regulator of lineage specification during differentiation of spinal-cord neurospheres. To functionally validate this finding, we quantified OLIG2-positive (oligodendrocytic) and GFAP-positive (astrocytic) cells by IF after growth-factor withdrawal (Fig. 5M,N). Consistent with the omics data, YAP1 knockout cultures showed a significant increase in OLIG2⁺ cells, while the proportion of GFAP⁺ cells fell sharply.

### Activation of Hippo/YAP1 signaling in spinal cord stem cells reduces oligodendrocytic gene expression while increasing ependymal cell genes and blocking proliferation

Findings from the loss-of-function experiments prompted us to examine the opposite scenario—gain of YAP1 function—by expressing a constitutively active YAP1 mutant (5SA) in neurospheres^46^. This construct increased total YAP1 while reducing its inactive phosphorylated form (pYAP) (Fig. 6A). We then performed transcriptomic and proteomic profiling under both proliferative and differentiation conditions.

**Figure 6.**
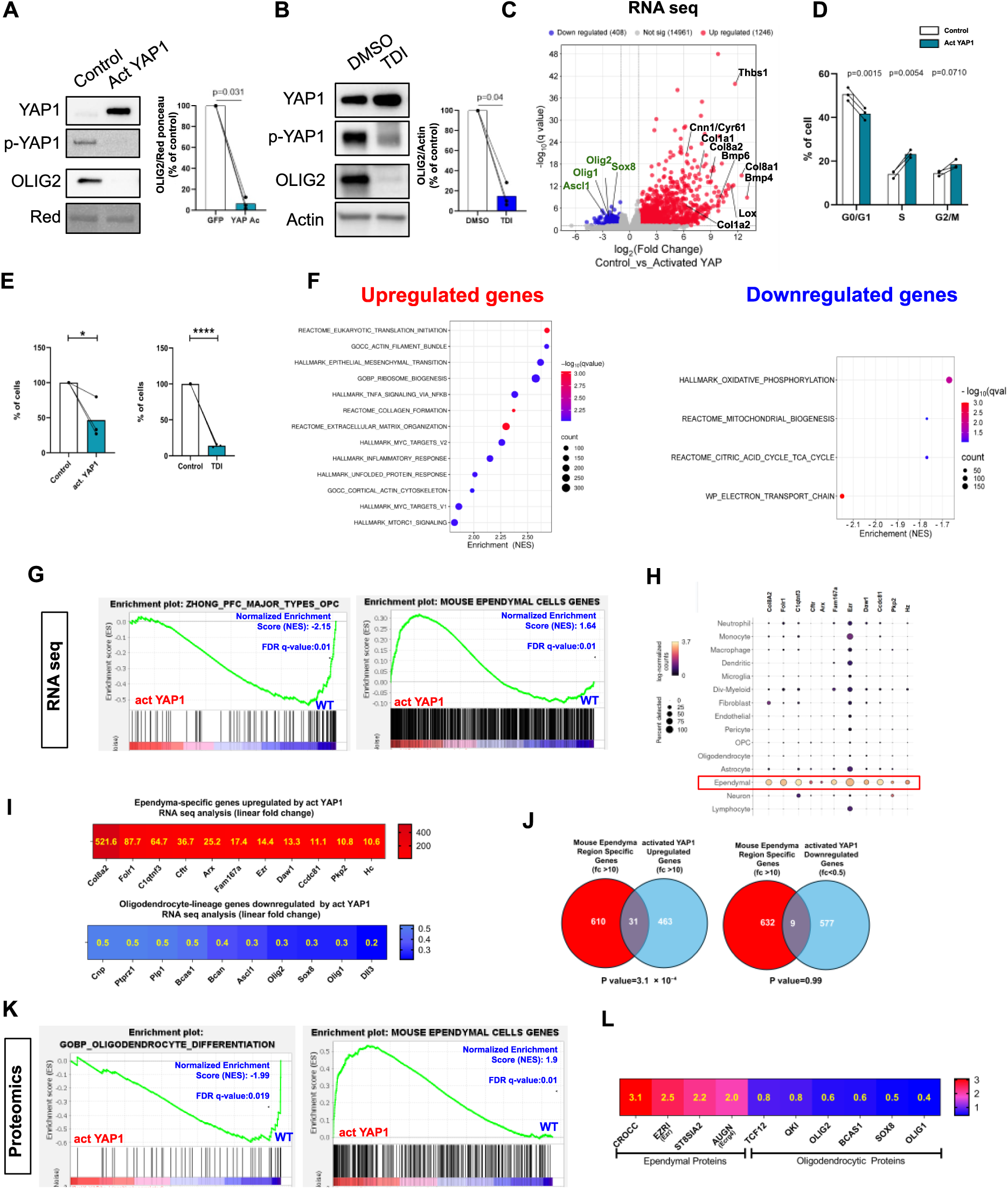
**Impact of 5SA YAP1 overexpression in proliferating neurospheres.** A. Western blot of total YAP1, phosphorylated YAP1 (p-YAP1), and OLIG2 in proliferative neurospheres transduced with control (GFP) or constitutively active YAP1 (5SA) viruses. Ponceau Red was used as a loading control. Right: Quantification of OLIG2 normalized to Ponceau and expressed relative to control virus. Active YAP1 significantly reduces OLIG2 expression (paired t-test, p value shown, n = 3). B. Western blot of YAP1, p-YAP1, and OLIG2 in proliferative neurospheres treated with DMSO or the YAP1 inhibitor TDI (5 µM). Actin was used as a loading control. Right: Quantification of OLIG2 normalized to Actin and expressed relative to DMSO-treated cells. TDI treatment significantly reduces OLIG2 expression (paired t-test, p value shown, n = 3). C. Volcano plot of bulk RNA-seq comparing control (GFP) and YAP1-activated transduced cells. Differentially expressed genes (|log₂ (fold change)| > 1, q < 0.05) are shown. Canonical oligodendrocyte lineage genes (green, left) are significantly downregulated, whereas extracellular matrix (ECM) components and BMP pathway genes (black, right) are strongly upregulated in YAP1-activated cells. D. Cell cycle analysis by flow cytometry (propidium iodide) in control (GFP, white) and YAP1-activated (blue) transduced cells. Bar graphs show the percentage of cells in G0/G1, S, and G2/M phases. Each dot represents an independent experiment (n = 3). Activated YAP1 decreases G0/G1 and increases S phase (paired t-test). E. Effect of 5SA YAP1 overexpression (left) or TDI treatment (5 µM, right) on cell number after 7 days of culture. Values expressed as percentage of control (GFP vector or DMSO). Mean ± SEM, n=3 independent experiments. Paired t-test: *p<0.05, p<0.0001. F. Bubble plots of enriched or depleted pathways (Table S3) in activated YAP1 vs GFP transduced cells. G. GSEA of RNA-seq from proliferative spinal cord neurospheres transduced with GFP or activated YAP1 viruses. YAP1 enhances ependymal signatures and reduces OPC signatures. NES and FDR q-values are shown. H. Dot plot of ependymal-specific genes strongly upregulated by activated YAP1 (depicted in heatmap I), adapted from the adult spinal cord single-cell atlas (Milich et al., J Exp Med, 2021). Dot size indicates the proportion of cells expressing each gene, and color represents average expression (yellow = high, purple = low). The ependymal population, highlighted in red, shows marked specificity. I. Heatmaps of selected genes with significant (p < 0.05) expression changes. Top: 11 ependymal-specific genes (see panel H) strongly upregulated by activated YAP1 (fold change >10). Bottom: 10 oligodendrocyte-lineage genes downregulated (fold change <0.5). Fold changes (Act YAP1 vs. GFP) are shown in yellow within heatmap cells. J. Venn diagrams showing the overlap between mouse ependyma-specific genes (fold change >10; Table S4, derived from Table S1 Ghazale et al., Stem Cell Reports, 2019) and genes either upregulated (fold change >10, left) or downregulated (fold change <0.5, right) in neurospheres transduced with activated YAP1 virus. Upregulated genes display strong enrichment (hypergeometric test, P = 3.1 × 10⁻⁴), whereas downregulated genes do not (p = 0.99), indicating that YAP1 activation promotes ependyml-specific gene expression. K. GSEA of cell-type-specific signatures in proteomic datasets from proliferative spinal cord neurospheres transduced with GFP or activated YAP1 viruses. Plots show enrichment of ependymal cell signatures in YAP1-activated cells and depletion of oligodendrocyte differentiation signatures. Normalized Enrichment Scores (NES) and False Discovery Rate (FDR) q-values are indicated. L. Heatmap of representative differentially expressed proteins (p < 0.05) between activated-YAP1 and GFP-transduced proliferative neurospheres. Oligodendrocyte-lineage proteins are shown in red/purple, astrocytic-lineage proteins in blue. Yellow numbers indicate linear fold change for each protein.

Overexpression of 5SA YAP1 triggered a strong transcriptional response, with 1,246 genes upregulated and 408 downregulated (|log₂ fold change| > 1, q-value<0.05. Fig. 6C; Table S3). GSEA/ORA analyses highlighted activation of protein synthesis, actin remodeling, and ECM programs, including collagens (*Col1a1, Col1a2, Col8a1, Col8a2*) and *Thbs1* (Fig. 6C, F). Several oncogenic signatures were also enriched (MYC targets, EMT, inflammatory and TNFα/NF-κB signaling), while mitochondrial and metabolic pathways—including oxidative phosphorylation, the TCA cycle, and electron transport—were repressed (Fig. 6F).

These changes suggested effects on proliferation. To test this, we analyzed cell cycle distribution and found that 5SA YAP1 expression significantly increased the proportion of cells in S and G2/M phases, with a corresponding reduction in G0/G1 (Fig. 6D). Paradoxically, however, total cell numbers after 7 days were markedly reduced in 5SA YAP1–transduced cultures (Fig. 6E). To investigate this further, we used TDI, a pharmacological activator of YAP1 that inhibits the upstream kinase LATS1/2^47^. TDI reduced pYAP levels (Fig. 6B), promoted nuclear YAP1 translocation in a subset of neurosphere cells (Fig. S6G), and, consistent with 5SA overexpression, also led to a pronounced decrease in cell number (Fig. 6E).

5SA YAP1 overexpression also profoundly affected lineage-specific programs. Oligodendrocyte-lineage genes—including the key transcription factors *Ascl1*, *Olig1*, *Olig2*, and *Sox8*—were strongly downregulated, as shown by the volcano plot (Fig. 6C) and GSEA (Fig. 6G). The heatmap in Fig. 6I highlights 10 representative genes suppressed under these conditions. OLIG2 reduction was confirmed at the protein level by WB in 5SA YAP1– overexpressing and TDI-treated cells (Fig. 6A,B). Conversely, YAP1 activation induced robust upregulation of ependymal-cell markers. GSEA revealed enrichment for mouse spinal cord ependymal signatures (Table S3, Fig. 6G), with dramatic induction (10- to 520-fold) of genes such as *Arx, C1qtnf3, Ccdc81, Cftr, Col8a2, Daw1, Ezr, Fam167a, Folr1, Hz*, and *Pkp2*, all highly enriched in ependymal cells according to single-cell atlases (Fig. 6H,I). Venn analysis confirmed significant overlap, with 31 upregulated genes ≥10-fold mapping to ependymal-specific signatures (Fig. 6J; Table S4), whereas downregulated genes showed no such enrichment.

Proteomic profiling mirrored the RNA-seq results: 5SA YAP1 overexpression elevated proteins linked to actin/ECM remodelling, translation, and cell-cycle progression, while diminishing those involved in mitochondrial oxidative phosphorylation and lipid β-oxidation (Table S3). Globally, oligodendrocyte-lineage proteins were depleted and ependymal markers enriched (Fig. 6K). The heatmap in Fig. 6L underscores this shift, showing some down-regulated oligodendrocyte proteins alongside four up-regulated ependymal-specific proteins highlighted by single-cell gene spinal cord atlas (Fig. S8): CROCC (Rootletin, ciliary rootlet component), EZR (Ezrin, membrane–actin linker), ST8SIA2 (NCAM polysialyltransferase), and AUGN/*Ecrg4*, a WNT-signalling modulator^48^.

Collectively, these findings demonstrate that YAP1 activation profoundly reprograms spinal cord stem cells, repressing oligodendrocytic programs while inducing ependymal-cell signatures and reducing cell numbers.

### Activation of Hippo/YAP1 blocks glial differentiation and promotes ependymal identity

Given that YAP1 activation profoundly remodels the transcriptomic and proteomic landscapes of proliferative spinal-cord neurospheres, we next asked whether these cells could still differentiate after growth-factor withdrawal (Fig. 7). Under these conditions, RNA-seq revealed 10,903 up- and 3,243 down-regulated genes—an even greater shift than in proliferating cultures (Fig. 7B). GSEA showed strong repression of astrocytic and oligodendrocytic programs and suppression of mitochondrial pathways (Fig. 7D,E). Consistently, GFAP and OLIG2 were nearly undetectable by WB and IF (Fig. 7A,J). In contrast, GSEA and the volcano plot indicated enrichment of cell cycle–associated gene sets, consistent with robust PCNA expression detected by WB in transduced cells (Fig. 7A,C–E). This was further supported by cell-cycle profiling, which revealed increased DNA synthesis without a corresponding rise in the G2/M population, suggesting that while 5SA-YAP1 promotes replication, cells stall before division (Fig. 7F).

**Figure 7.**
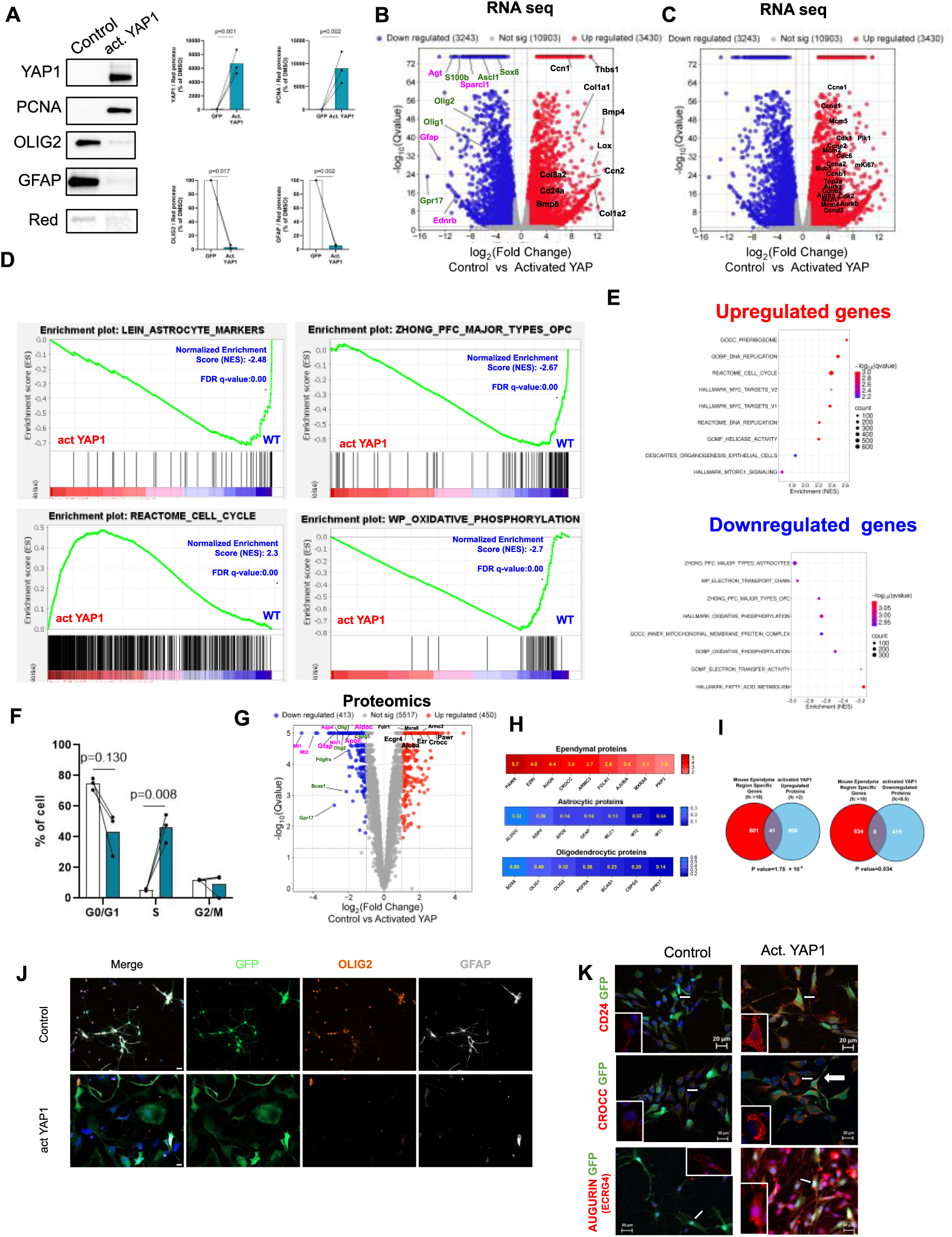
**Effect of constitutively active 5SA YAP1 on neurosphere differentiation.** For this experiment, proliferative neurospheres were transduced with either GFP or constitutively active (5SA) YAP1-expressing virus, then differentiated for 3 days by growth factor withdrawal on laminin. A. Western blot of YAP1, PCNA, OLIG2, and GFAP in differentiated neurospheres (Ponceau Red = loading control). Bar graphs (right) show protein levels normalized to Ponceau Red, expressed as % of GFP control. 5SA YAP1 overexpression increases YAP1 and PCNA, while reducing OLIG2 and GFAP. Data: n = 3, paired t-test. B–C. Volcano plots of bulk RNA-seq comparing GFP control and 5SA YAP1-transduced cells (|log_2_(fold change)| > 1 ; q < 0.05). B. Oligodendrocyte (green) and astrocyte (pink) lineage genes are downregulated, while ECM-related and BMP pathway genes (black) are upregulated in YAP1-activated cells. C. Key cell cycle regulators are upregulated, consistent with enhanced cell cycle signaling. D. GSEA of RNA-seq from differentiated spinal cord neurospheres transduced with GFP or activated YAP1. Plots show downregulation of astrocyte, OPC, and oxidative phosphorylation genes, with upregulation of cell cycle genes, consistent with a shift toward proliferation and reduced differentiation/oxidative metabolism. NES and FDR q-values are indicated. E. Bubble charts of selected pathways from Table S3 showing significant enrichment or depletion in GFP control vs. YAP1-activated cells. F. Cell cycle analysis by flow cytometry (propidium iodide) in control (GFP, white) and YAP1-activated (blue) cells. Bar graphs show the percentage of cells in G0/G1, S, and G2/M phases. Each dot = independent experiment (n = 3). Activated YAP1 tends to reduce G0/G1 (p = 0.130) and significantly increases S phase (p = 0.008), with no change in G2/M (paired t-tests). G. Volcano plot of proteomics comparing GFP control and YAP1-activated differentiated neurosphere cells (|log₂fold change| > 1; q < 0.05). Oligodendrocyte (green) and astrocyte (pink) lineage proteins are downregulated, while ependymal-specific proteins (black) are enriched among upregulated proteins. H. Heatmap of selected differentially expressed proteins (p < 0.05) in differentiated neurospheres (YAP1-activated vs. GFP). Markers of ependymal cells (red), astrocytes (blue, middle), and oligodendrocytes (blue, bottom) are shown. Linear fold change values are indicated in yellow. I. Venn diagrams showing overlap between region-specific mouse ependymal genes (fold change >10; Table S4, derived from Table S1 Ghazale et al., Stem Cell Reports, 2019) and proteins upregulated (fold change >2, left, left) or downregulated (fold change <0.5 right) in differentiated neurospheres expressing activated YAP1. Significant enrichment is observed in both sets (hypergeometric test: up, p = 1.75 × 10⁻⁹; down, p = 0.034), stronger in the upregulated set, indicating that YAP1 primarily promotes ependymal-specific protein expression. J. Triple immunofluorescence for GFAP, GFP, and OLIG2 in differentiated neurospheres transduced with GFP control or activated YAP1. YAP1 activation strongly suppresses GFAP and OLIG2 expression. Scale: 10 µm. K. Immunofluorescence for the ependymal proteins CROCC and AUGURIN (AUGN/Ecrg4) in differentiated neurospheres transduced with GFP control or activated YAP1. Insets show magnified views of cells indicated by white arrows. YAP1 activation strongly increases CROCC and AUGURIN expression. Scale: 20 µm. Nuclei, DAPI (blue), all photographs.

Proteomics analysis reinforced this picture: proteins characteristic of oligodendrocytes and astrocytes dropped, while those encoded by ependymal-cell specific genes rose (Fig. 7G,H). Venn diagram analyses show that YAP1-upregulated proteins overlapped significantly with ependymal-specific genes (41 shared; p = 1.75 × 10⁻⁹), whereas the down-regulated set showed only a weak association (p = 0.034) (Fig. 7I). Collectively, these data indicate that sustained YAP1 activity suppresses glial differentiation while promoting an ependymal-like transcriptional program. To corroborate these findings at the cellular level, we performed IF for CROCC and AUGN/*Ecrg4* —two novel ependymal markers identified in this study-—and for CD24a, a well-established ependymal marker previously reported by us and others^49,50^. Constitutive activation of YAP1 using the S5A variant markedly increased the expression of all three proteins (Fig. 7K), further supporting the role of YAP1 signaling in promoting the acquisition of an ependymal identity.

## Discussion

A subset of ependymal cells lining the adult mouse spinal cord central canal retains bona fide neural stem cell properties, with long-term self-renewal in vitro and multipotent differentiation capacity^1,2,4,6–8,22^. Although their chromatin landscape is primed for oligodendrocyte fate^25^, these cells predominantly generate astrocytes after injury^22^. The molecular signals that bias their lineage choice remain poorly understood. To address this gap, we investigated how OPCs emerge in vitro from neurosphere cultures derived from the adult spinal cord.

This study led to several findings. Our data show that spinal-cord neurospheres already express important oligodendrocyte-lineage transcription factors (OLIG1/2, SOX4, NKX2.2, TCF4) with the notable exception of SOX10, implying a poised oligodendrocytic-differentiation program that is readily activated in culture. Because SOX10 expression depends on neuronal^51^ and endocrine cues—particularly thyroid hormones^52^ —its absence suggests that additional extrinsic signals must be supplied in vitro to fully unlock the oligodendrocyte program. Neurosphere cells also express SOX9 and NFIA, pivotal drivers of astrocytic fate^31^. In previous work^9^, we showed that these cells express radial-glial markers (GLAST and FABP7/BLBP), indicating that spinal-cord stem cells adopt a radial glia-like phenotype in vitro. By co-expressing both astrocytic and oligodendrocytic transcription factors, these cells appear to exist in an hybrid state, poised to differentiate toward either the astrocytic or oligodendrocytic lineage. Upon adhesion of neurospheres and growth factor withdrawal, the expression of oligodendrocyte-lineage transcription factors is broadly reduced, yet remains selectively maintained in OPCs, underscoring their distinct molecular identity within the culture. This coexistence of dual, lineage-primed transcriptional programs—followed by bifurcation upon fate commitment—parallels observations made in other stem cell systems, most notably in hematopoietic progenitors^53^.

Leveraging *Pdgfra*^H2B-GFP^ cultures, we captured newly formed PDGFRA⁺ OPCs in neurospheres for molecular characterization. Transcriptomic and proteomic profiling confirmed oligodendrocyte-lineage commitment, yet this commitment remained plastic: a subset of PDGFRA-GFP⁺ cells can still form new neurospheres and produced GFP⁻ progeny. This echoes prior work showing BMP-driven reprogramming of OPCs into stem-like cells^54^. In line with this, our proteomics detected endogenous BMP1 (Table S1), suggesting a possible intrinsic trigger for reversibility.

Molecular profiling of these newly-generated OPCs highlighted a pronounced up-regulation of chromatin remodelers consistent with an epigenetically driven fate switch. Concurrently, these cells upregulated glutamatergic neurotransmitter receptors together with some synaptic scaffold genes, implying that they are already primed for neuron-glia communication as reported in the brain^55^. Signalling pathways classically linked to astrocytic specification were concurrently repressed. Notch-regulated transcription factors dropped sharply, and EGFR levels declined at both mRNA and protein levels—an outcome that may stem from asymmetric receptor segregation previously reported in subventricular-zone neural stem cells^56^. These nascent OPC also down-regulated Tenascin-C (TNC), an extracellular-matrix ligand that potentiates EGFR/Ras signalling^57^. Given that EGFR activation drives astrocyte differentiation^58^, the coordinated silencing of EGFR and its amplifier TNC, together with diminished Notch input, is likely to tip lineage commitment toward oligodendrogenesis at the expense of astrogenesis.

A central outcome of this study is that Hippo/YAP1 signaling regulates lineage specification between oligodendrocytes and astrocytes and modulates ependymal-cell identity and cilia-associated gene expression in cultured spinal-cord neural stem cells. Building on our earlier RNA-seq observations of Hippo/YAP1 target enrichment in mouse and human ependymal cells^15^, we validated robust YAP1 and TEAD1 protein expression in ependyma across species. Loss- and gain-of-function experiments in neurospheres then revealed distinct effects on proliferation and differentiation.

With respect to proliferation, YAP1 loss caused only minor transcriptomic and proteomic changes, and we detected no obvious impact on neurosphere growth. This is consistent with the work by Huang et al., who reported normal growth of *Nestin*-Cre Yap1-null neurospheres from embryonic brain^59^. This suggests YAP1 is largely inactive in proliferating stem cells or compensated by its paralogue TAZ. Conversely, overexpression of the constitutively active 5SA-YAP1 variant strongly upregulated proliferation-associated genes, potently blocked differentiation by suppressing both astrocytic and oligodendrocytic markers, and increased the proportion of cells in S phase. However, this was not sufficient to promote entry into G2/M or to increase the total cell number. TDI treatment reproduced these effects, supporting an antiproliferative role for Hippo/YAP1 activation in this context. YAP1 can act as either a pro-or anti-proliferative factor depending on context^60,61^, and in our system it seems to adopt the latter role—perhaps priming cells for mitosis while still requiring additional cues to complete division. This interpretation is consistent with the high levels of YAP1 we observed in spinal ependymal cells in vivo, which are quiescent under normal conditions but re-enter the cell cycle rapidly following spinal cord injury.

YAP1’s influence on lineage choice becomes apparent when growth factors are withdrawn to trigger differentiation. Under these conditions, RNA-seq, proteomics, and IF all show a sharp decline in astrocytic markers in YAP1-KO neurospheres compared with wt cultures. This echoes the work of Huang et al., who reported defective astrocyte formation in cortical neurospheres lacking YAP1 and traced the defect to disrupted crosstalk between Hippo/YAP and BMP signaling—BMP being a key astrocytic driver^59^. A similar breakdown in Hippo-BMP synergy likely underlies the impaired differentiation we observe in the spinal-cord context. More unexpectedly, our study uncovered a novel role for YAP1 in regulating the ependymal cell phenotype. Integrated RNA-seq and proteomic analyses demonstrate that YAP1 depletion markedly down-regulates several ependymal-specific markers, whereas YAP1 overexpression conversely up-regulates a subset of ependymal-specific genes. A particularly striking example is *Col8a2*, a gene highly specific to ependymal cells in spinal cord single-cell atlases (Fig. S7) whose expression decreased nearly fivefold in YAP1 KO (Fig. 5E) and increased over 500-fold upon Hippo/YAP1 activation (Fig. 6C and I). In vivo, dual YAP/TAZ deletion produces hydrocephalus and depletes ependymal cells^62^, and CNS-specific *Yap1* ablation likewise blocks their generation^63^. Together, our in vitro and these in vivo observations strongly suggest that YAP1 as a pivotal regulator of ependymal-cell formation and maintenance.

Ependymal cells are characterized by the presence of cilia, and we found that a substantial fraction of the ependymal genes regulated by YAP1 encode cilia-related proteins. One notable example is *Crocc* (also known as Rootletin), a gene highly enriched in ependymal cells (Fig. S8) that encodes a structural protein responsible for anchoring cilia to the cytoskeleton^64^, thereby ensuring their mechanical stability in ciliated cells. Crocc protein was reduced in *Yap1* KO cells (Fig. 5K) and increased upon YAP1 overexpression (Fig. 6L, 7H, K). This downregulation of cilia-associated genes in *Yap1*-deficient cells is possibly due to reduced expression of transcription factors that orchestrate ciliogenesis, such as *FoxJ1* and *Rfx2*, both of which were significantly downregulated in the absence of YAP1. The involvement of YAP1 in ciliogenesis has recently emerged in other biological contexts^65,66^, and our findings further support this emerging role.

Finally, we revealed an important link between YAP1 and oligodendrogenesis: (i) YAP1/TEAD decline during OPC formation (Fig. 4); (ii) YAP1 loss upregulates oligodendrocyte markers, notably *Olig1/2* key transcription factors (Fig. 5); and (iii) activated YAP1 or TDI suppresses OPC markers (Figs. 6–7). *Olig2*—one master regulator of oligodendrocyte fate— is increased upon YAP1 loss and nearly abolished by YAP1 activation, positioning YAP1 as a potent repressor of *Olig2* and the oligodendrocyte program. Beyond transcriptional effects, YAP1 may act epigenetically: during OPC formation we observed upregulation of chromatin-remodeling genes, and YAP1 loss further increased key modifiers (*Ezh2, Nsd1/2, Kmt2c, Setd2*) (Table S2), suggesting YAP1 helps shape chromatin accessibility to restrain oligodendrocyte differentiation.

A limitation of our study is that Hippo/YAP1’s role was explored mainly in vitro. After spinal injury, ependymal cells predominantly yield astrocytes rather than oligodendrocytes. Testing whether YAP1 loss in vivo enhances oligodendrogenesis would therefore be highly informative.

In conclusion, our work provides a detailed molecular characterization of spinal cord stem cells and their capacity to generate OPCs. It identifies Hippo/YAP1 as a regulator of ependymal identity, ciliogenesis, and lineage choice between astrocytes and oligodendrocytes. The RNA and proteomic datasets generated here constitute a valuable resource for future studies, and the enrichment of previously unrecognized genes and proteins in OPC opens new avenues for dissecting lineage specification. Since ependymal cells with stem-like features persist in the human spinal cord^3,11,27^, modulating YAP1 activity may hold translational potential for promoting repair in traumatic and degenerative spinal cord injury.

## Supporting information

Table S1 RNA seq and Proteomics Analyses of PDGFRA-GFP+ vs GFP-cells

Table S2 KO YAP RNA seq and Proteomics Analyses

Table S3 RNA seq and proteomics activated YAP1

Table S4 Mouse and human ependymal cell-specific genes.

Table S5 List of 91 genes of figure 5F

Table S6 Antibody list

## Acknowledgements

We gratefully acknowledge financial support from the following organizations: Wings for Life Spinal Cord Research Foundation (WFL, Austria); Institut pour la Recherche sur la Moelle Épinière et l’Encéphale (IRME, France), AFM-Téléthon (Association Française contre les Myopathies, AFM, France), ARSEP (Fondation pour l’Aide à la Recherche sur la Sclérose en Plaques, France), International Foundation for Research in Paraplegia (IRP, Swiss). S.E.J, R.C., J.A.C, were supported by ARSEP, AFM and Wings for Life fellowships respectively. We thank Dr. Muriel Perron, Dr. J. Wrana, and Dr. S. Meilhac for generously providing the *Yap1*^fl/fl^ mouse line and for their expert advice. We are also grateful to Dr. B. Oumesmar and Dr. P. Soriano for providing the *Pdgfra*^H2B-GFP^ KI mouse line. Our work also relied on the outstanding support of the PPM, MRI, MGX, RAM, and RHEM core facilities. The Orbitrap Exploris 480 mass spectrometer used at the Montpellier Proteomics Platform (PPM, BioCampus) for proteomics analysis was co-financed by the European Regional Development Fund (ERDF) and the Occitanie region.

## Authors contribution

J.-P.H. conceived and supervised the project; S.E.J., R.C., L.Z, M.C and J.P.H. analyzed the data. S.E.J. and R.C. performed most of the experiments, with assistance from J.A.C., S.L. L.G., A.N., S.H., C.R., H.G., and V.H.P. G.P., F.V.F., and L.B. were responsible for obtaining patient consent and collecting human spinal cord samples (Fig. 4E). E.L. and C.R. provided a custom-made antibody targeting AUGN. S.U., M.S., and K.E.K. carried out the proteomics experiments and subsequent data analysis. S.E.J. drafted the initial version of the manuscript and prepared the first set of figures. J.P.H. wrote the main text, finalized all figures, and prepared the submitted version. All authors read and approved the final manuscript.

## Materials and methods

### Animals

Animals were housed in animal care facilities in compliance with the Committee of the National Institute of Health and Medical Research (INSERM) and in accordance with the European Council directive (2010/63/UE). All animals had diet and water *ad libitum* and were socially housed in a 12h light-dark cycle. Adult C57Bl/6, *Yap1*^fl/fl^ ^41,42^ and *Pdgfra*-^H2B-GFP29^ mice (3–4 months) were euthanized by CO2 before spinal cord extraction for culture or histology.

### Cell Cultures

Spinal cord neural stem cell cultures were established from adult mouse spinal cords following a previously described protocol^67^. The base medium consisted of DMEM/F12 (Gibco), 1× N2 supplement (Gibco), 0.25× B27 supplement (Gibco), 2 mM L-glutamine (Gibco), 2 mg/L ciprofloxacin (Sigma), and 10 mg/L gentamicin (Gibco). For proliferation, cells were seeded at a density of 4,000 cells/cm² onto PolyHEMA-coated culture supports (Sigma) in base medium further supplemented with 2 mg/L Fungin (InvivoGen), 2 mg/L heparin (Sigma), and 10 μg/L each of EGF and FGF2 (Peprotech). Differentiation was induced by plating 50,000 cells/cm² onto surfaces pre-coated with poly-D-lysine and laminin (1 μg/cm²; Sigma) in base medium lacking EGF, bFGF2, and heparin. All experiments were performed on neurospheres maintained for fewer than 10 passages after initial derivation from adult spinal cord tissue.

### Proliferation assays and cell counting

Cell proliferation was assessed by EdU incorporation (Fig. 1E and 5C). Cells were incubated with 10 μM 5-ethynyl-2′-deoxyuridine (EdU) for 2 hours, followed by detection using the method described in Salic and Mitchison^68^. To directly measure the effect of YAP1 overexpression and TDI treatment on cell number, cells were seeded in six wells per condition of a 24-well plate, and total cell counts were determined after two weeks using a Z2 automated cell counter (Beckman Coulter).

### Pharmacological and Genetic Manipulation of YAP1

Pharmacological activation of the Hippo/YAP1 pathway was performed by adding 5 μM TDI-011536 (MedChemExpress) to the culture medium on days 0 and 3. To induce expression of a constitutively active YAP1 mutant (YAP1-5SA^46^), in which all five canonical LATS1/2 phosphorylation sites (Ser61, Ser109, Ser127, Ser164, and Ser381) are mutated to alanine, cells were transduced with a recombinant adenovirus encoding GFP–YAP1-5SA (VectorBuilder). Control cells were transduced with an adenovirus encoding GFP alone.

YAP1 Knockout Experiments: To eliminate YAP1 expression in cultures derived from Yap^fl/fl^ mice^41^ and wild-type littermates, neurospheres were transduced with either a GFP–Cre– expressing adenovirus (Ad5-GFP-Cre) or a GFP-only control virus. GFP⁺ cells were isolated by FACS (BD Melody) and expanded for downstream analyses, including RNA sequencing and proteomic profiling. In *Yap1*^fl/fl^ cultures, Cre recombinase excised exon 2 of the YAP1 gene, introducing a frameshift mutation that abolished full-length protein production^41^.

### Immunostainings and quantification

Cells cultured on glass coverslips were fixed with 4% paraformaldehyde (PFA) for 15 minutes and stored in PBS containing 0.05% sodium azide (Sigma). Antibodies used for immunofluorescence are listed in Table S6. Briefly, cells were permeabilized and blocked in a buffer containing 0.3% Triton X-100 and 5% donkey serum, followed by incubation with primary antibodies overnight at 4 °C. Secondary antibodies (Jackson ImmunoResearch) were applied for 1 hour at room temperature, followed by nuclear staining with DAPI (1 μg/mL) and mounting with Permount (Invitrogen). Images were acquired using a Zeiss AxioImager microscope equipped with an Apotome module and analyzed using Zeiss or Fiji software. Most cell quantifications were performed on digitized images, and positive cells were counted using ImageJ software. EGFR, TNC, and HES1 stainings shown in Figure 4A were directly evaluated and quantified under the microscope. For each condition, at least 100 cells were counted across 2 to 3 coverslips, with cells distributed over a minimum of 10 randomly selected fields. Immunofluorescence on mouse spinal cords (Fig. 4D) was performed on animals intracardially-perfused with PFA 4%. The immunohistochemistry shown in Fig. 4E was carried out on a human spinal cord sample from a 53-year-old donor, using the protocol established in a prior publication from our laboratory^3^.

### RNA profiling

Cells were collected and RNA was extracted using the RNeasy Kit (QIAGEN). RNA profiling of PDGFRA-GFP⁺/GFP⁻ and YAP1 knockout cells was performed at the MGX platform (IGF, France), while RNA profiling of cells transduced with constitutively active YAP1 (5SA) was conducted by BGI genomics (China and Poland).

At the MGX facilities, total RNA were quantified on Fragment Analyzer (Agilent Technologies, Santa Clara, CA, USA) using the standard sensitivity RNA kit (DNF-471-0500). Libraries were prepared using the SMART-Seq Stranded Kit (Takara Bio) according to the manufacturer’s instructions. The size distribution of the resulting libraries was assesed using a Fragment Analyzer (Agilent Technologies, Santa Clara, CA, USA) and the libraries were quantified using the KAPA Library quantification kit (Roche, Basel, Switzerland). Sequencing was performed on a NovaSeq 6000 (Illumina, San Diego, CA, USA) using the single-read 1×100 nt protocol on one lane of an SP flow cell. Demultiplexing and FASTQ files generation were carried out using Illumina’s bcl2fastq software (v2.20.0.422). The quality control of raw data and demultiplexed reads was assessed using respectively Illumina’s Sequencing Analysis Viewer (SAV) software and the FastQC (v0.11.8) software from the Babraham Institute. Contaminant screening was performed using FastQ Screen (v0.14.0, Babraham Institute). A splice junction mapper, TopHat (v2.1.1) [1] (using Bowtie v2.3.5.1 [2]), was used to align the reads to the *Mus musculus* genome (mm39 C57BL6J) with a set of gene model annotations (gff file downloaded from UCSC on 2021-02-08). Final read alignments having more than 6 mismatches were discarded. Samtools (v1.9) was used to sort and index the alignment files. Then, gene counts were obtained using FeatureCounts (v2.0.0) [3]. As the data is from a strand-specific assay, the reads have to be mapped to the opposite strand of the gene (-s 2 option). Before statistical analysis, genes with less than 15 reads (cumulating all the analyzed samples) were filtered out. Differentially expressed genes were identified using the Bioconductor [4] package DESeq2 (v1.26.0, R v3.6.1) [5]. Data were normalized using the DESeq2 normalization method. Genes with adjusted p-value below 5% (according to the FDR method from Benjamini-Hochberg) were called differentially expressed.

secondary structure, and mRNA is enriched by oligo (dT) attached magnetic beads. After reacting at a suitable temperature for a fixed time period, RNAs are fragmented with fragmentation reagents. Then first-strand cDNA is generated using random hexamer-primed reverse transcription with the fragmented RNA as template. To synthesize the second-strand cDNA, the synthesis reaction system is prepared and dUTP is used to replace dTTP. After obtaining the double strand cDNA product, it is converted to blunt end with end repair reaction. After cDNA end repairment, a single ‘A’ nucleotide is added to the 3’ ends of the blunt fragments through A tailing reaction. Then the library adapters are ligated to the two ends of the cDNA with ligation reaction. Finally, the library products are amplified through PCR reaction and subjected to quality control. Next, the single-stranded library products are produced via denaturation. The reaction system for circularization is set up to get the single-stranded circularized DNA products. Any single stranded linear DNA molecules will be digested. The final single strand circularized library is amplified with phi29 and rolling circle amplification (RCA) to make DNA nanoball (DNB) which carries more than 300 copies of the initial single stranded circularized library molecule. The DNBs are loaded into the patterned nanoarray and sequencing reads with PE 150 bases length are generated on DNBSEQ-T7 platform generating 20M paired reads per sample. Raw sequencing data were filtered using SOAPnuke (v1.5.2), a quality control software independently developed by BGI. The following parameters were used: -l 15, -q 0.2, and -n 0.05.

RNA-seq data were analyzed using the DESeq2 package. Volcano plots and bubble charts were generated using the SRPlot web tool^69^ (https://www.bioinformatics.com.cn). Functional enrichment and cell phenotype analyses were performed using Gene Set Enrichment Analysis (GSEA) software^32^ and Over-Representation Analysis (ORA)^35^ via the Enrichr web platform^70^.

### Proteomic profiling

Cells were lysed in a buffer containing 5% SDS and 100mM of TEABC (Triethylammonium bicarbonate). Protein digestion was performed on S-Trap micro columns (Protifi) following the manufacturer’s instructions. Obtained peptides were labelled with TMT (Thermo Fisher) and pooled. The labelled peptide samples were fractionated into eight fractions using high pH reversed-phase kit (Pierce). Fractions were analysed with nanoLC MSMS using an Exploris 480 equipped with FAIMS source. Raw spectra were analysed with MaxQuant (v 2.0.3.0) using UniProt Reference proteome of Mus musculus (Proteome ID UP000000589 ; https://www.uniprot.org/). Statistical analyses were performed using Perseus (v1.6.15.0)^71^. Functional enrichment and cell phenotype analyses were performed using GSEA, ORA analyses and 1D annotation enrichment Perseus^72^. Volcano plots and bubble charts were generated using the SRPlot web tool^69^.

### Cell cycle analysis by FACS

Cell cycle analysis was performed with a Propidium Iodide (PI) assay. Cells were resuspended in 70% EtOH in PBS prior to an incubation with 0.25mg/ml of RNAse-A DNase-free for 30 minutes. Staining was performed 10 minutes before the analysis with 50 ug/ul of PI. Acquisitions were realized with the Miltenyi MACSQuant and analysis were completed on FlowJo.

### Western Blot

Cells were lysed in RIPA buffer (Sigma) supplemented with protease and phosphatase inhibitor cocktails (complete™ ULTRA Tablets and PhosSTOP™, Roche). Protein concentrations were quantified, and 10 µg of total protein per sample was mixed with Laemmli sample buffer containing β-mercaptoethanol. Samples were denatured by heating at 95 °C for 5 minutes and loaded onto Mini-PROTEAN® TGX Stain-Free™ Gels (Bio-Rad). Proteins were separated by SDS-PAGE and transferred to PVDF membranes using the Trans-Blot® Turbo Transfer System (Bio-Rad). Membranes were blocked and incubated with primary and secondary antibodies as detailed in Table 1. Signal detection was carried out using the Clarity™ Western ECL substrate (Bio-Rad), and chemiluminescence was captured with a ChemiDoc™ Imaging System (Bio-Rad). Band intensities were quantified using Image Lab™ software (Bio-Rad).

### Statistical analysis

Data analysis was performed using Microsoft Excel (Version 2508), GraphPad Prism (v10), Perseus (v1.6.15.0)^71^, and Cytoscape (v3.10.2)^34^. Data were assumed to follow a normal distribution, although this was not formally tested. Statistical analyses were conducted using the tests indicated in each figure. Exact p-values were reported directly on the graphs or defined as follows: p < 0.05 (*), p < 0.01 (**), and p < 0.001 (***).

### Data Availability

Bulk RNA-seq datasets have been deposited in the NCBI Gene Expression Omnibus (GEO) under the following accession numbers: GSE274710 for the PDGFRA-GFP+ vs GFP-cell data, GSE274111 for the YAP1 KO cells and GSE275944 for the constitutively active YAP1 cells. Mass spectrometry proteomics data have been deposited in the ProteomeXchange Consortium via the PRIDE partner repository. For PDGFRA-GFP+/GFP-cells, proteomics datasets are available under the identifiers PXD055075 (proliferation condition) and PXD055072 (differentiation condition). Note that one sample in PXD055072 were excluded from analysis due to a FACS sorting issue, which resulted in insufficient enrichment of GFP cells in the GFP⁺ fraction. For the YAP1 KO vs wt cells, proteomics data are available under the dataset identifier PXD055079. For the constitutively active YAP1 cells, datasets are available under the identifiers PXD055110 (proliferation) and PXD055106 (differentiation).

### Language Editing Assistance

Stylistic improvements and clarity enhancements were assisted by ChatGPT (OpenAI), an artificial intelligence–based language model.

## Declarations

The authors declare no conflicts of interest.

## Ethical approval

This study was performed in line with the principles of the Declaration of Helsinki. Approval was granted to Dr Luc Bauchet by the French “Agence de la biomédecine”, n° SPGED19-3-4692, April, 2nd, 2019 for human spinal cord samples. Informed consent was obtained from relatives of organ donors included in the study (Fig. 4E).

---

**Figure S1.**
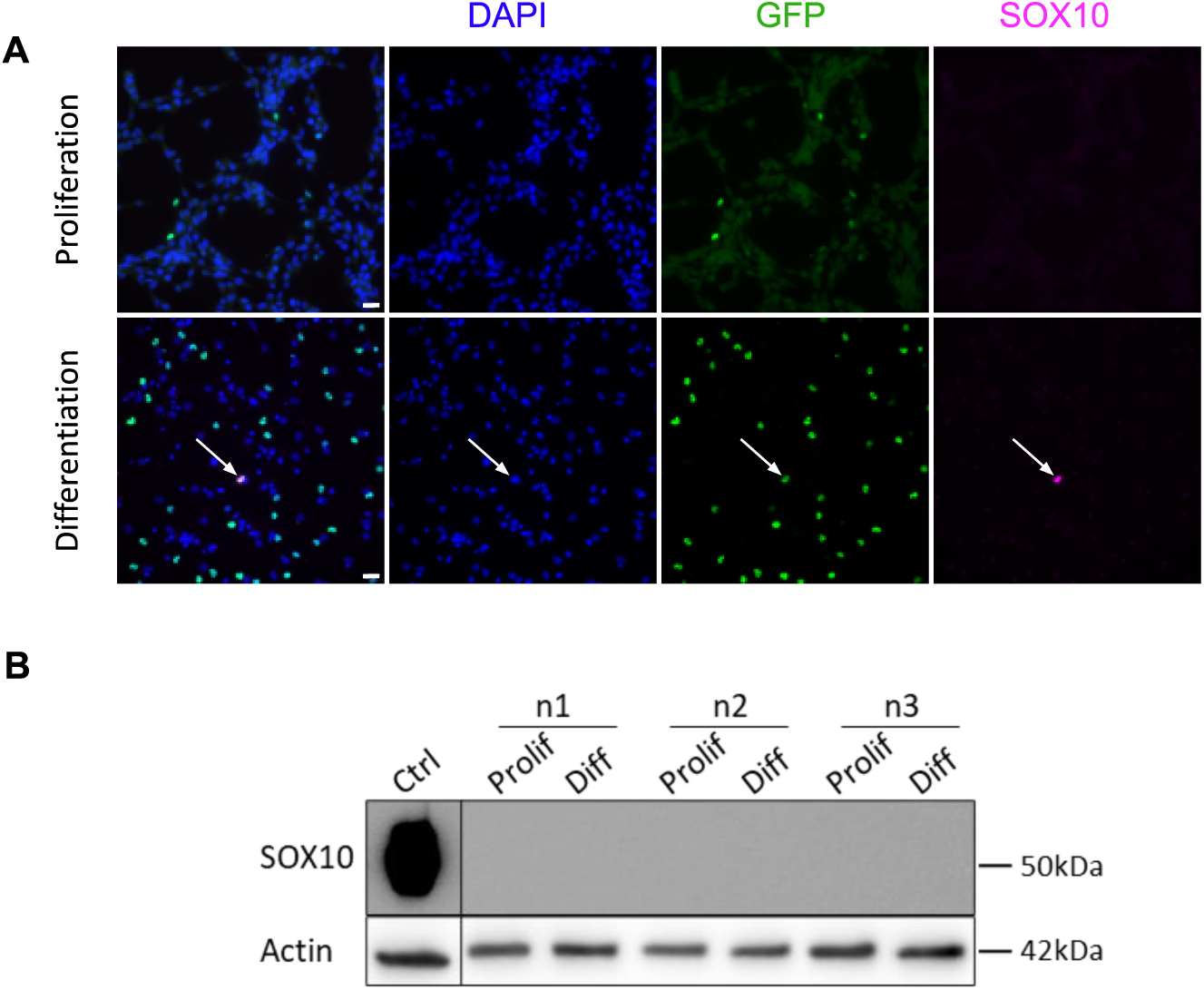
Lack of Sox10 expression in adult spinal-cord neurospheres (related to. figure 1) A. Immunofluorescence of neurospheres derived from adult PDGFRA-^H2B-GFP^ spinal cord mice in proliferation (top) or after 5 days of differentiation (bottom). GFP (green), SOX10 (magenta), and nuclei (DAPI, blue). Arrows indicate rare GFP⁺ SOX10⁺ cells after differentiation. Scale: 20 µm. B. WB for SOX10 in three independent neurosphere preparations (n1–n3) under proliferative (Prolif) or differentiating (Diff) conditions. Proteins extracted from the BT138 human oligodendroglioma cell line serve as positive control (Ctrl). Actin is the loading control. No SOX10 band is detected in neurosphere samples, confirming negligible protein expression.

**Figure S2.**
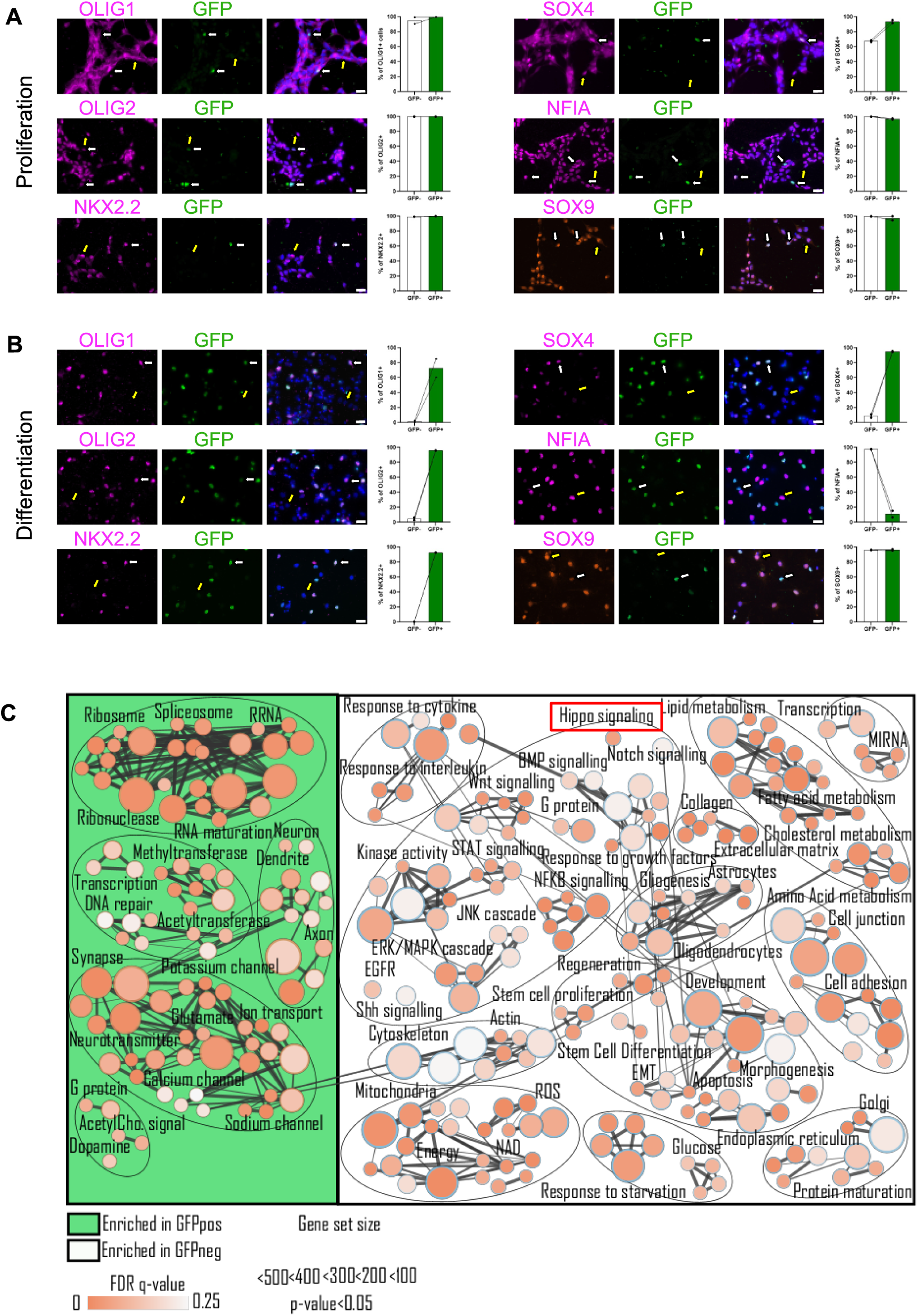
Transcription factor expression and pathway enrichment in PDGFRA-GFP⁺ and GFP⁻ spinal cord neurosphere cells (related to. Fig. 1). A–B Immunofluorescence transcription factors factor (OLIG1, OLIG2, NKX2.2, SOX4) and astrocytic TFs (NFIA, SOX9) in GFP⁺ and GFP⁻ cells under proliferative (A) and differentiation (B) conditions. Representative images (left) and quantification (right, n = 2) are shown. In proliferation, both GFP⁺ and GFP⁻ cells express similar levels of all six TFs. Upon differentiation, GFP⁻ cells downregulate oligodendroglial TFs, whereas GFP⁺ cells maintain their expression along with SOX9 but show reduced NFIA. White arrows: GFP⁺ cells; yellow arrows: GFP⁻ cells. Scale: 20 µm. C Global RNA analysis. Bubble plot (Cytoscape 3.3.0; Genome Res. 2003, 13:2498) summarizing gene-set enrichment of RNA-seq data from sorted GFP⁺ versus GFP⁻ cells (FDR-adjusted p < 0.05). Bubble size = gene-set size; color intensity = adjusted p-value. GFP⁺ cells (green background, left) are enriched in ribosomal, spliceosomal, and RNA-processing modules. GFP⁻ cells (white background, right) show enrichment in Hippo, NF-κB, and metabolic pathways linked to astrocytic specification. The Hippo pathway (red box) is selectively enriched in GFP⁻ cells.

**Figure S3.**
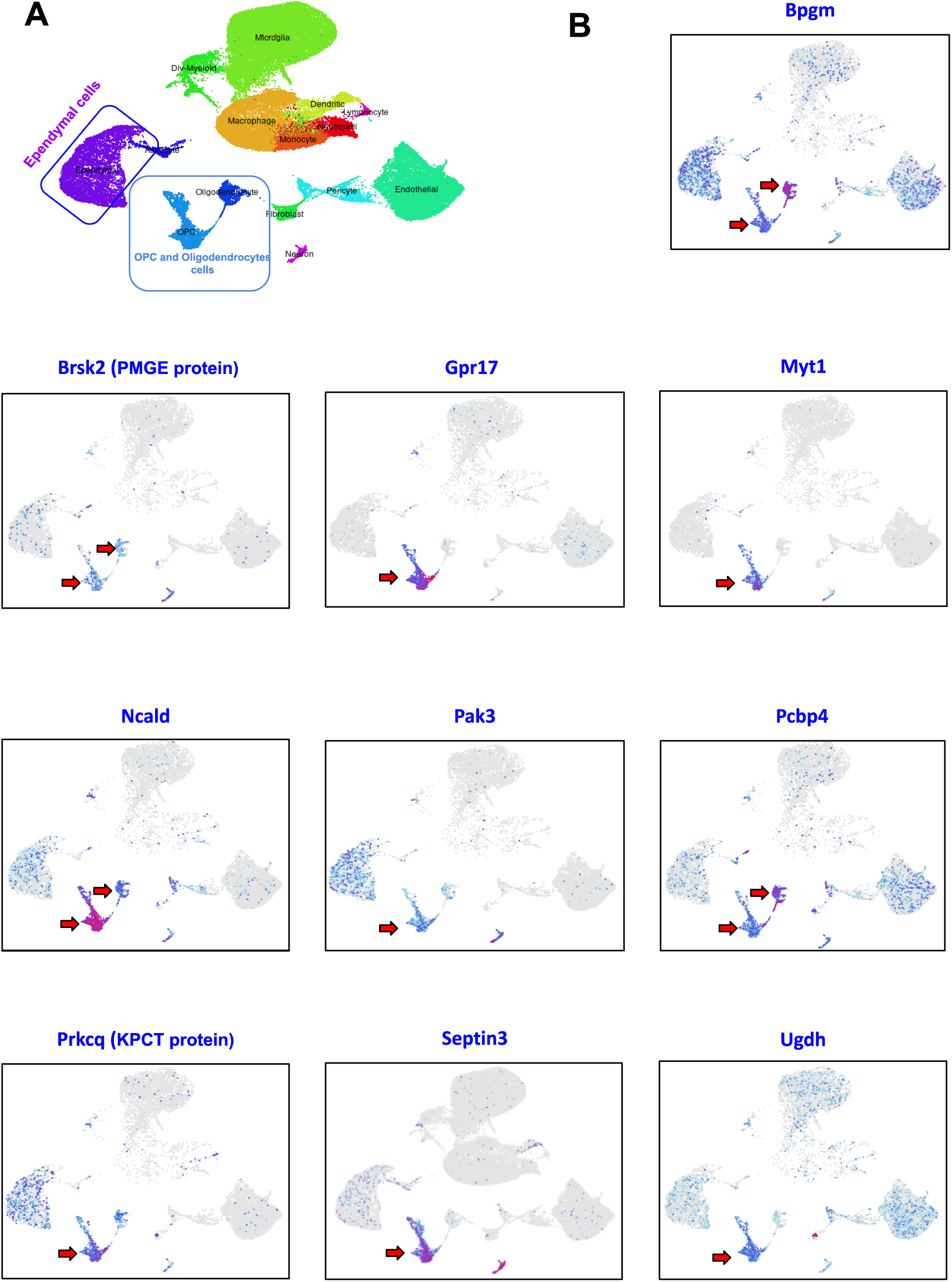
Single-cell atlas–based expression patterns of 10 genes encoding proteins enriched in PDGFRA-GFP⁺ vs. GFP⁻ cells from proliferating spinal cord neurospheres (related to. Figure 3H) A. UMAP of spinal cord cells grouped by cell type based on transcriptomic profiles (Milich et al., 2021; PMID: 34132743; https://jaeleelab.shinyapps.io/sci_singlecell/). Distinct clusters correspond to microglia, macrophages, neutrophils, monocytes, lymphocytes, OPCs, oligodendrocytes, astrocytes, ependymal cells, neurons, fibroblasts, pericytes, and endothelial cells. B. UMAP plots showing expression of the indicated genes. Levels are color-coded from grey (not detected) to blue (low) and red (high). Red arrows highlight enrichment in OPCs and/or oligodendrocytes. Nine of ten genes are predominantly expressed in these lineages, except Pak3. Ependymal and astrocyte cells show little or no expression, supporting the lineage specificity of proteins identified in GFP^+^ cells (Fig. 3H).

**Figure S4.**
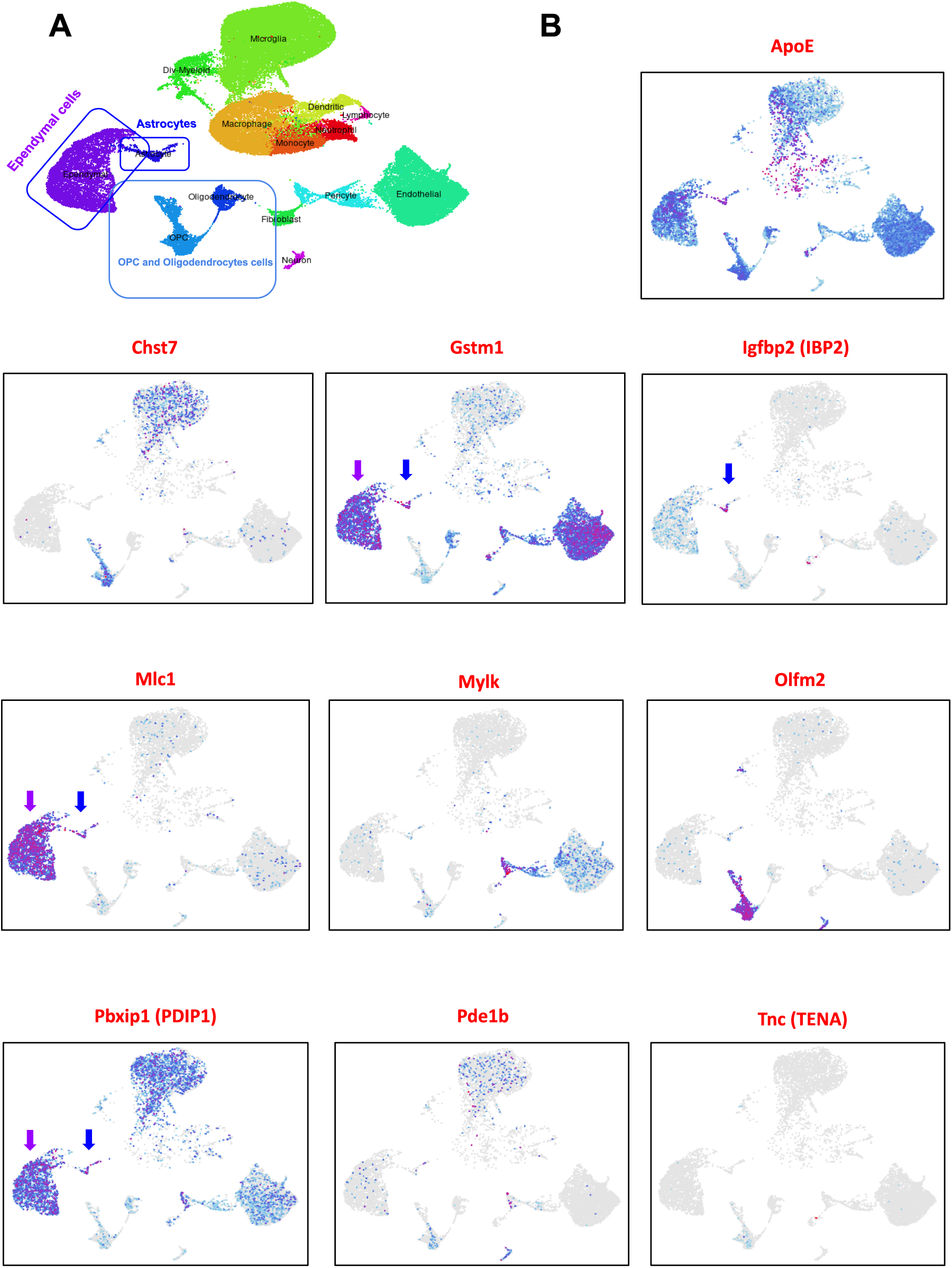
Single-cell atlas–based expression patterns of 10 genes encoding proteins enriched in GFP⁻ vs. PDGFRA-GFP+ cells from proliferating spinal cord neurospheres (related to. Figure 3H) A. UMAP of spinal cord cells grouped by cell type based on transcriptomic profiles (Milich et al., 2021; PMID: 34132743; https://jaeleelab.shinyapps.io/sci_singlecell/). Distinct clusters correspond to microglia, macrophages, neutrophils, monocytes, lymphocytes, OPCs, oligodendrocytes, astrocytes, ependymal cells, neurons, fibroblasts, pericytes, and endothelial cells. B. UMAP plots showing the expression of the indicated genes. Expression levels are depicted using a color scale ranging from grey (not detected) to blue (low expression) and red (high expression). Purples and blue arrows indicate enrichment in ependymal cells and astrocytes respectively. Four out of the ten genes analyzed are predominantly expressed in ependymal or astrocytes.

**Figure S5.**
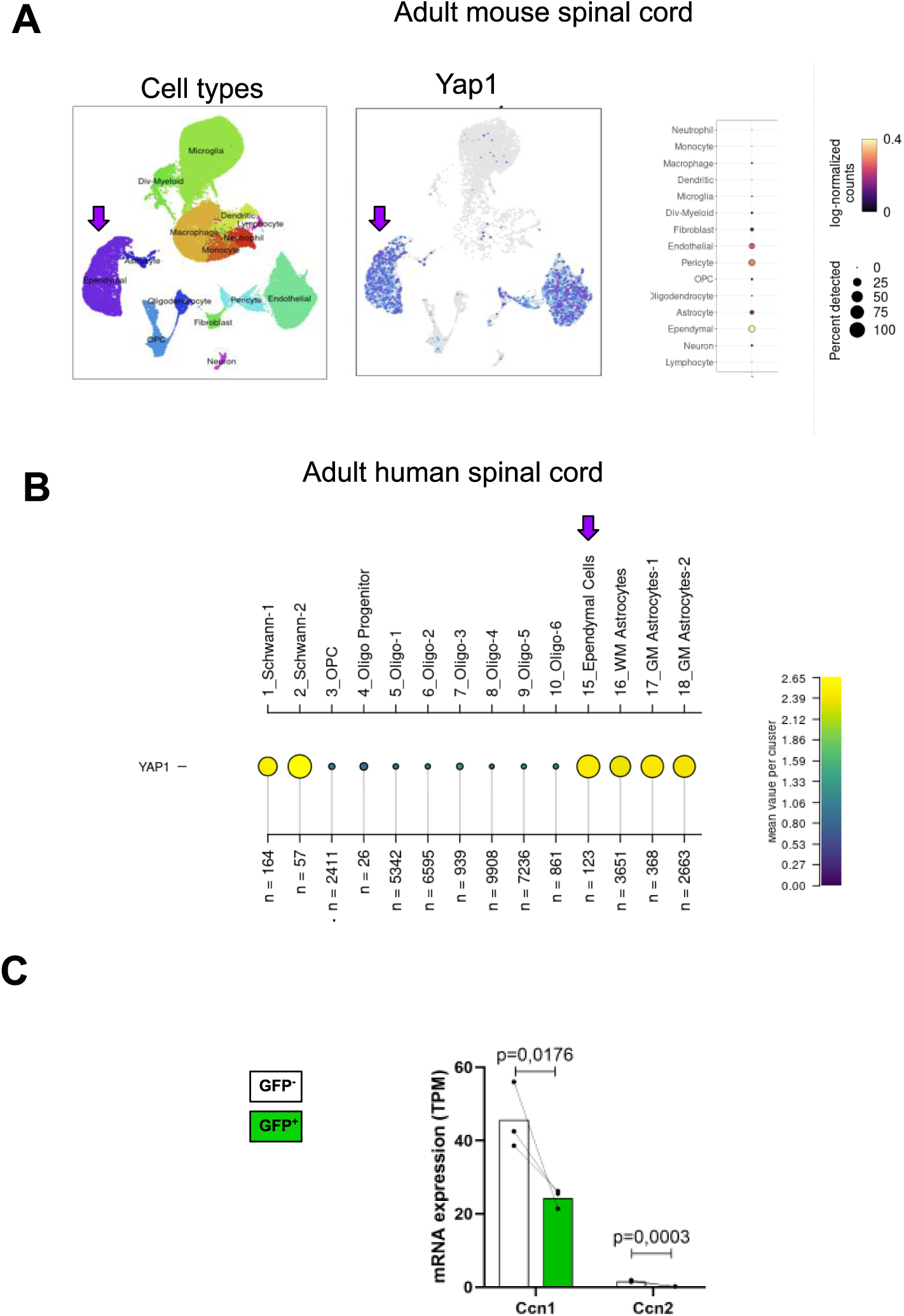
Yap1 expression across adult mouse and human spinal cord cell types, and differential expression of *Ccn1/Ccn2* in PDGFRA-GFP⁺ and GFP⁻ cells (related to. **figure 4).** A. Yap1 expression across adult mouse spinal cord cell types. Left: UMAP of spinal cord cells grouped by cell type based on transcriptomic profiles (Milich et al., 2021; PMID: 34132743; https://jaeleelab.shinyapps.io/sci_singlecell/). Middle: UMAP plots showing Yap1 expression enriched in ependymal cells, pericytes, and endothelial cells. Right: Dot plot (adapted from dataset from Milich et al., 2021) showing Yap1 expression across major cell types, with marked specificity in ependymal cells compared to astrocytes, OPCs, and oligodendrocytes. Dot size = proportion of expressing cells; dot color = average expression (yellow, high; purple, low). Purple arrows highlight ependymal cells. B. Yap1 expression across adult human spinal cord cell types. Dot plot from the single-cell RNA-seq atlas (Yadav et al., Neuron 2023; 111:328–344.e7) showing expression level and proportion of YAP1⁺ cells across annotated clusters. Dot size = % of cells expressing YAP1; color = average expression (log-normalized counts). n = number of cells per cluster. Yap1 is mainly expressed in ependymal cells (cluster 15) and in white and gray matter astrocytes, with lower expression in oligodendrocyte lineage cells (clusters 3–10). Data from https://vmenon.shinyapps.io/humanspinalcord/. C. Bar graphs of mean mRNA levels (RNA-seq, n = 3) for *Ccn1 (Cyr61)* and *Ccn2 (Ctgf)*, canonical Hippo/YAP1 targets, in FACS-sorted spinal cord PDGFRA-GFP⁺ (green) and GFP⁻ (white) cells. Unpaired t-test.

**Figure S6.**
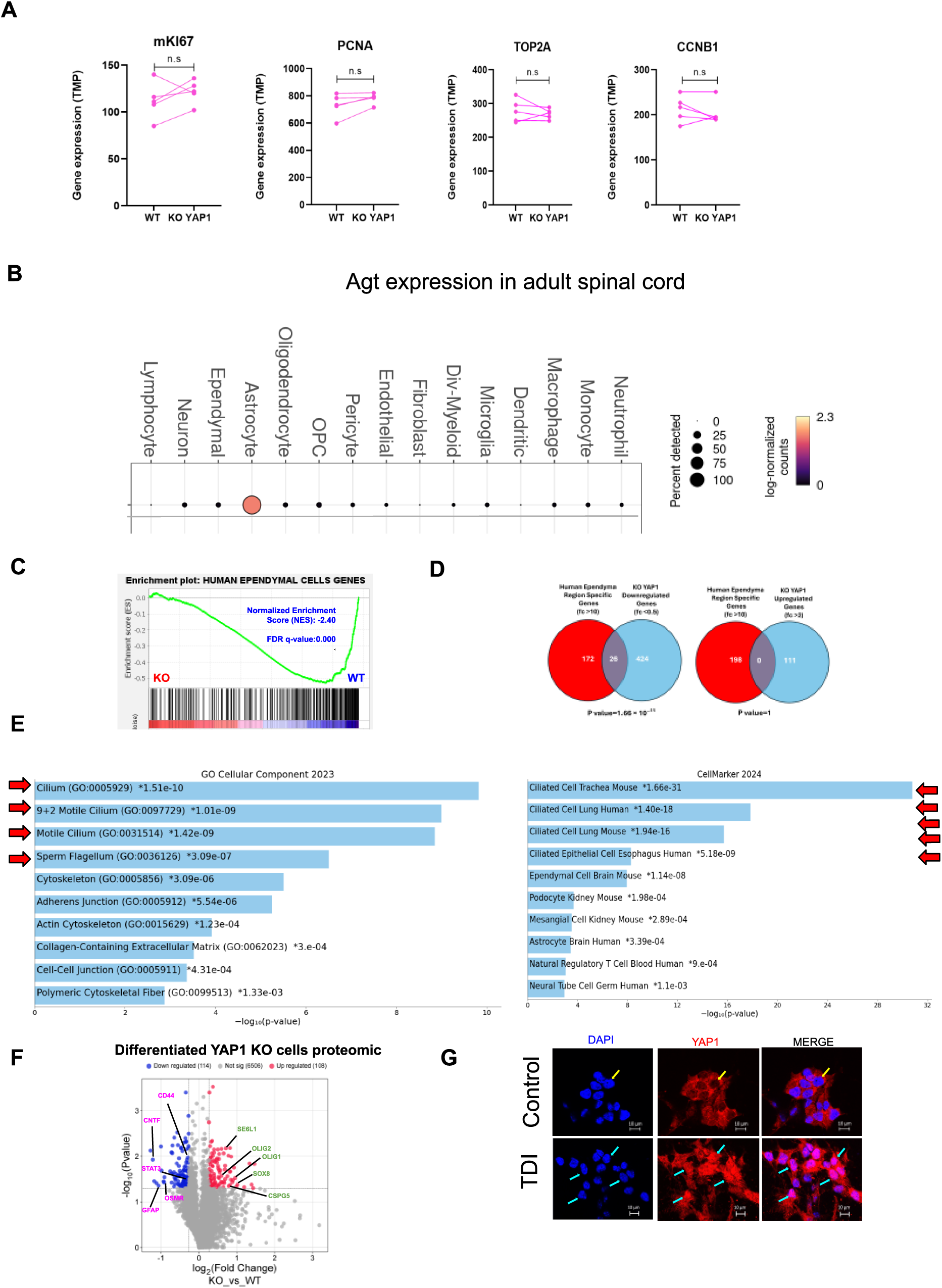
Further analysis of gene and protein expression alterations in YAP1 knockout neurospheres (related to. Figure 5). A. Paired dot plots of *Mki67*, *Pcna*,*Top2a*, and *Ccnd1* transcript levels (TPM, RNA-seq; n = 5) in WT and YAP1 knockout (KO) neurospheres. Lines connect paired samples. No significant differences (n.s., unpaired t-test). B. Agt expression across adult mouse spinal cord cell types. Dot plot from a single-cell RNA-seq dataset (Milich et al., 2021; PMID: 34132743, https://jaeleelab.shinyapps.io/sci_singlecell/) showing *Agt* (angiotensinogen) expression. Dot size = % of cells per cluster; dot color = average expression (log-normalized counts). Agt is predominantly expressed in astrocytes, with little or no expression in other cell types. Data from C. GSEA of the human ependymal cell signature (Table S4) using differentially expressed genes from RNA-seq of WT and YAP1 KO differentiated neurospheres (Table S2). Plots show significant depletion of ependymal genes in YAP1 KO cells. NES and FDR q-values are indicated. D. Venn diagrams showing the overlap between human ependymal genes (fc >10; gene list Table S4) derived from Table S2 in Ghazale et al., *Stem Cell Reports*, 2019) and genes either downregulated (fc <0.5, left) or upregulated (fc >2, right) in differentiated YAP1 KO neurospheres. Strong enrichment is observed among downregulated genes (hypergeometric test, p = 1.66 × 10⁻¹¹), but not among upregulated ones (p = 1), indicating that YAP1 loss reduces ependymal gene expression.. E. Gene enrichment analysis of the 91 genes identified in Fig. 5H that are (i) enriched in mouse ependymal cells and (ii) downregulated in differentiated YAP1 KO neurospheres. Left: GO Cellular Component 2023 analysis, with bars ranked by –log₁₀(p). Downregulated genes are strongly enriched for cilium-related and cytoskeletal structures (“Cilium,” “Motile Cilium,” “Adherens Junction”), indicating loss of cilia programs after YAP1 loss. Right: Enrichr analysis (CellMarker database) highlights signatures of ciliated epithelial cells (tracheal, lung) and brain ependymal cells. Numbers indicate adjusted p-values (asterisk). Together, these enrichments show that YAP1 sustains ependymal/ciliated-cell identity in spinal cord stem cell cultures. F. Volcano plot of proteomic profiling comparing GFP⁺ and GFP⁻ cells under differentiative conditions. This plot is related to Figure 5E Proteomics Volcano plot. Statistical significance was determined using a p-value threshold < 0.05 (n=5), rather than a q-value in Fig. 5E. Differentially expressed proteins were identified using a fold-change>1.2 (i.e. |log_2_(fold-change|>0.26). Astrocyte-specific proteins are highlighted in pink and oligodendrocyte lineage proteins in green. G. IF for YAP1 (red) in neurospheres maintained with growth factors and treated with TDI. Top: DMSO control; bottom: 5 µM TDI, 1 h. Nuclei counterstained with DAPI (blue). In controls, YAP1 is mostly cytoplasmic and excluded from the nucleus (yellow arrows), whereas TDI induces nuclear accumulation (blue arrows), with many cells showing both nuclear and cytoplasmic staining. Scale: 10 µm.

**Figure S7.**
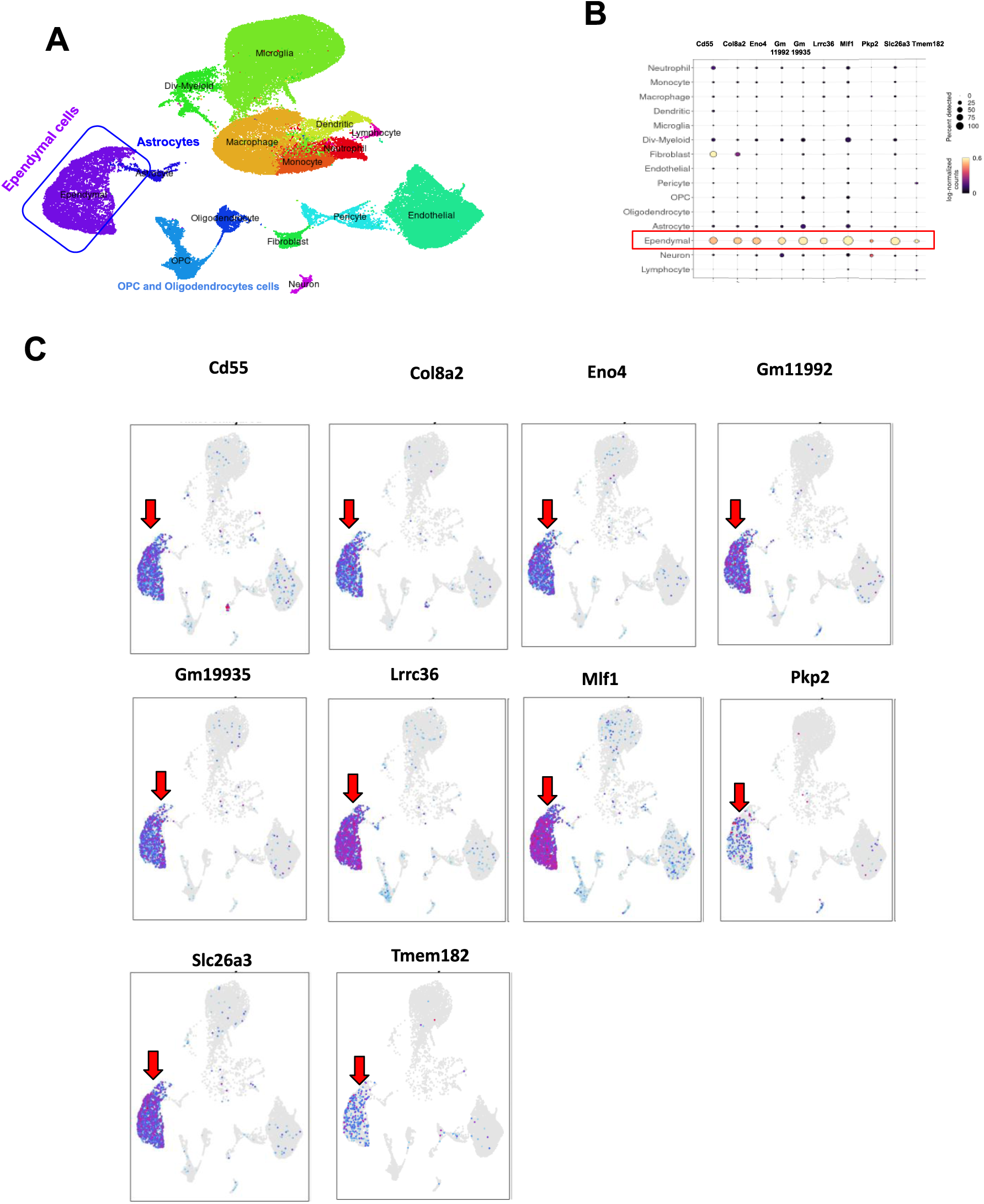
Expression profiles of nine genes significantly down-regulated in YAP1-KO neurospheres under differentiation conditions, mapped onto a single-cell spinal-cord atlas (related to. Fig. 5F). A. UMAP of spinal cord cells grouped by cell type based on transcriptomic profiles (Milich et al., 2021; PMID: 34132743; https://jaeleelab.shinyapps.io/sci_singlecell/). Distinct clusters correspond to microglia, macrophages, neutrophils, monocytes, lymphocytes, OPCs, oligodendrocytes, astrocytes, ependymal cells, neurons, fibroblasts, pericytes, and endothelial cells. B. Dot plot adapted from dataset in Milich et al., 2021 (PMID: 34132743; https://jaeleelab.shinyapps.io/sci_singlecell/) showing expression of nine genes across major spinal cord cell types. Expression is highly specific to ependymal cells (red arrow). Dot size = proportion of cells expressing the gene; dot color = average expression (log-normalized counts; yellow = high, purple = low). Cell types are listed on the y-axis. C. UMAP plots showing expression of the indicated genes. Expression ranges from grey (not detected) to blue (low) and red (high). Red arrows highlight specific expression of the nine genes in ependymal cells.

**Figure S8.**
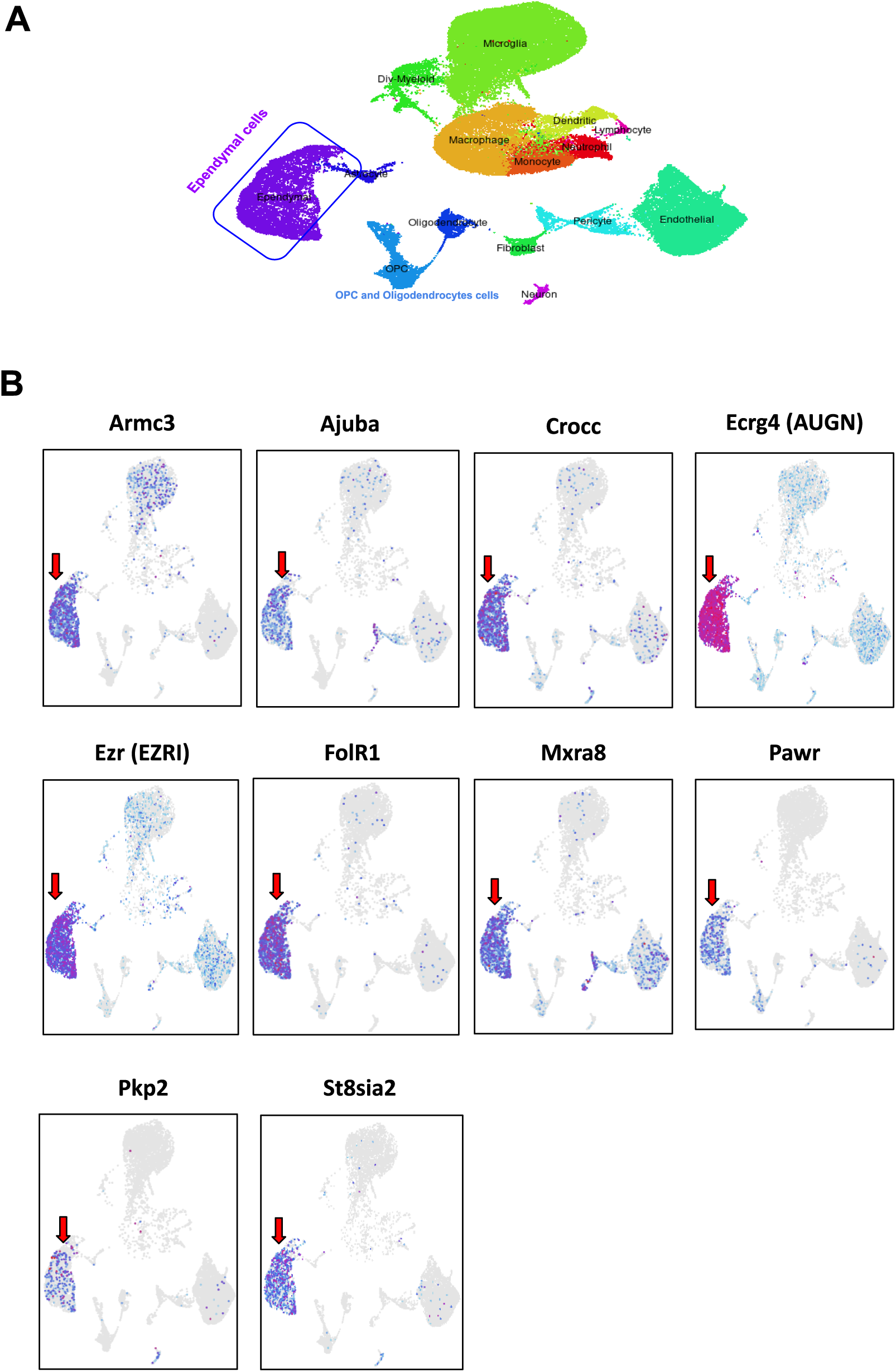
Single-cell atlas–based expression patterns of 10 genes encoding proteins enriched in differentiated spinal cord neurospheres overexpressing 5SA-YAP1 (related to. Figure 6L **and 7H).** A. UMAP of spinal cord cells grouped by cell type based on transcriptomic profiles (Milich et al., 2021; PMID: 34132743; https://jaeleelab.shinyapps.io/sci_singlecell/). Distinct clusters correspond to microglia, macrophages, neutrophils, monocytes, lymphocytes, OPCs, oligodendrocytes, astrocytes, ependymal cells, neurons, fibroblasts, pericytes, and endothelial cells. C. UMAP plots showing the expression of the indicated genes. Expression levels are depicted using a color scale ranging from grey (not detected) to blue (low expression) and red (high expression). Red arrows highlight enrichment of these 10 genes in ependymal cells.

**Table S1**. Transcriptomic and proteomic profiles of PDGFRA-GFP⁺ and GFP⁻ cells, including differential expression, functional enrichment, and cell phenotype analyses.

**Table S2:** Transcriptomic and proteomic profiles of YAP1 KO cells including differential expression, functional enrichment, and cell phenotype analyses.

**Table S3.** Transcriptomic and proteomic profiles of cells transduced with 5SA YAP1, including differential expression, functional enrichment, and cell phenotype analyses.

**Table S4.** List of human and mouse ependymal cell–specific genes used for GSEA and Venn diagram analyses in Figures 5F, 5H, 6G, 6J, 6K, and 7I (mouse gene list), as well as Figures S6C and S6D (human gene list). These gene sets were generated by selecting genes with at least a 10-fold higher expression in the ependymal region compared to the non-ependymal region. Both lists are derived from our previous publication: Table S1 and S2 in Ghazale et al., Stem Cell Reports, 2019 May 14; 12(5):1159–1177.

**Table S5.** List of the 91 genes that overlap between those down-regulated in YAP1-knockout differentiated neurospheres (Fig. 5H) and the ependymal-cell-specific signature identified in our earlier study (Ghazale et al., *Stem Cell Reports* 12: 1159–1177, 2019; see Table S1).

**Table S6.** Antibody list

## Notes

### Competing Interest Statement

The authors have declared no competing interest.

